# Benchmarking remote sensing methods to capture plant functional diversity from space

**DOI:** 10.1101/2025.10.22.683881

**Authors:** Javier Pacheco-Labrador, Ulisse Gomarasca, Daniel E. Pabon-Moreno, Wantong Li, Mirco Migliavacca, Martin Jung, Gregory Duveiller

## Abstract

The development of remote sensing methods to estimate plant functional diversity is limited by mismatches between ecology and remote sensing sampling schemes, and the limited representativeness of local field campaigns. The Biodiversity Observing System Simulation Experiment (BOSSE) provides a modeling framework for benchmarking new methodologies. We used BOSSE to simulate 180 different synthetic “Scenes” encompassing a two-year-long time series of plant trait maps and imagery of hyperspectral reflectance factors, spectral indices, sun-induced chlorophyll fluorescence, land surface temperature, and estimates of plant traits (optical traits). We used these simulations to answer five fundamental, yet unsolved, questions:

*Q1. How should remote sensing characterize functional diversity in large surfaces (sites)?* Diversity metric values saturate with the number of pixels involved, hampering comparisons between plant traits and remote sensing estimates in large areas. The average value of metrics computed over small samples should be used instead.

*Q2. Which sources of spectral information (or combinations thereof) can best capture plant functional diversity at the site scale?* Accounting for background effects is the key. Optical traits (remote sensing estimates of plant traits) are the best estimators for plant functional diversity. Other variables succeed when filtered out of the soil pixels; their combination did not yield additional advantages.

*Q3. How should remote sensing estimates be validated/compared with plant functional diversity measurements?* Leaf area index (LAI) is a better proxy of abundance than the pixel for *Q*_Rao,_ but not for variance-based partitioning. It is more sensitive to sample size, but also more resistant to suboptimal spatial resolution.

*Q4. When (in the phenological year) can remote sensing best capture site-scale plant functional diversity?* The estimation error decreased with LAI and stabilized at values above 1 m²/m².

*Q5. Which approaches and remote sensing variables are more resistant to the effects of suboptimal spatial resolution?* Optical traits, fluorescence, and reflectance factors were the most robust variables. Still, field data resolution needs to be degraded to match the sensor’s resolution. We found a relative spatial resolution threshold of ∼30 % (where the pixel is around three times larger than the plants).

Simulation frameworks like BOSSE enable testing methodologies beyond local contexts and address the current shortage of suitable global datasets, supporting the application and development of methods for assessing plant functional diversity with remote sensing. In the future, BOSSE could contribute to understanding observational results, refining and pre-testing new methodologies, and supporting the development of comparable experimental datasets.

## 1. INTRODUCTION

The development of remote sensing capabilities for studying plant diversity has increased significantly over the last decade (Torresani et al., 2024). The reasons behind this rise in interest stem from both the urgent need to develop a global biodiversity monitoring system (Gonzalez et al., 2023; Skidmore et al., 2015) and advancements in remote sensing technologies and sensors (Ma et al., 2020; Skidmore et al., 2021). As in other areas of environmental monitoring and research (e.g., climate change (Yang et al., 2013)), remote sensing could potentially play a key role as a provider of continuous and synoptic information for environmental protection, and in this specific case for the conservation of biodiversity (Cavender-Bares et al., 2022; Hansen et al., 2021) and ecosystem functioning (Abelleira Martínez et al., 2016; J. Cabello et al., 2018). However, despite the potential capabilities to capture different aspects of plant (and other taxa) diversity, contrasting results suggest that unresolved methodological issues limit advances in this field (Torresani et al., 2024).

The spectral variation hypothesis posits that the spatial heterogeneity captured by imagers is related to habitat and thus taxonomic richness (Palmer et al., 2002). Previous works have questioned this underlying hypothesis or its applicability. For example, simulations and experimental data from Fassnacht et al. (2022) showed that species identity, rather than taxonomic richness, governed spectral variability. Using simulations, Ludwig et al. (2024) refuted the notion that species richness drives spectral variability in grasslands and reported a strong dependency on spectral resolution. These works confirmed the results of earlier experiments, which had reported methodological (e.g., spatial resolution, metrics, preprocessing, or soil background effects) or contextual limitations (i.e., dependent on plant type or identity) to find the relationships predicted by the spectral variation hypothesis (Badourdine et al., 2023; Rossi et al., 2021a; Van Cleemput et al., 2023; Wang et al., 2018a, 2018b). The accumulated literature suggests that, beyond the intrinsic limitations to generalizing the spectral variation hypothesis (i.e., different species can be spectrally similar), numerous methodological challenges still hinder the full capability of remote sensing.

Beyond taxonomic diversity, remote sensing could capture functional diversity (i.e., the diversity of functional traits) thanks to the physical links between vegetation’s biophysical properties and spectral signals (Jetz et al., 2016). Such connections may serve as a means to access phylogenetic diversity (Schweiger et al., 2018). Remote sensing is already a key tool for monitoring vegetation properties across space and time (Homolová et al., 2013), which can be further advanced with the advent of new sensors and methods to monitor vegetation (Kattenborn et al., 2021; Zhang et al., 2021).

However, the estimation of plant functional diversity presents particularities and poses new challenges for remote sensing. Theoretical works (i.e., simulations using radiative transfer models) suggest that spectral diversity exhibits a strong inherent correlation with the diversity of vegetation properties (Pacheco-Labrador et al., 2022). Nevertheless, the same study shows that such relationships are not generalizable to all methods used to calculate functional diversity, functional diversity metrics (FDMs), or sensor features, and highlights gaps between remote sensing and ecology science that need to be overcome. Several studies have assessed the ability of different sensors and methods to capture plant functional diversity. The earliest works utilized airborne sensors to map plant functional traits and assess their diversity. In these works, plant trait measurements were used to build the statistical models estimating the trait maps (Asner et al., 2014; Schneider et al., 2017). Others compared the FDMs computed from drone or airborne imagery (reflectance factors and sun-induced chlorophyll fluorescence) with FMDs calculated from optical traits (remote sensing estimates of plant traits). These studies utilized ground data for validation (Tagliabue et al., 2020) or parameterization of radiative transfer models (Cimoli et al., 2024). Another line of work has empirically linked Sentinel-2 spaceborne imaging spectral reflectance factors with plant trait FDMs (Ma et al., 2019) or directly compared them with FDMs computed from reflectance and optical traits with those calculated from field-sampled plant traits (Pacheco-Labrador et al., 2022). However, even in these cases, field data were never sampled for validating remote sensing estimates of diversity. Worthy efforts in this direction were made by Hauser et al. (2021), who matched 3-by-3 sampling plot grids with Sentinel-2 pixels. Nonetheless, these remote sensing-dedicated approaches multiply the field sampling efforts, which must be species-based to be comparable with ecological studies. In addition, comparing the results of different studies is complex due to: 1) the numerous and non-overlapping varieties of sensors, sampling designs, diversity metrics, plant traits, and vegetation types encompassed in each research work; and 2) the limited spatial extents, since most often they happen in a single site. As a consequence, it is not possible to determine the limitations of each methodology and whether these can be applied globally. Furthermore, the gap between the variables often sampled by ecologists and remote sensing scientists (e.g., concentrations vs. contents) and the different effects these have on the spectral signals (Kattenborn et al., 2019) adds another layer of uncertainty to the exploitation of ecological datasets. Thus, while every study is valuable, the estimation of plant functional diversity from space still suffers from a lack of adequate datasets for validation of retrievals and benchmarking of methods. This makes it difficult to understand the requirements and limitations in terms of sensors, resolutions, diversity metrics, or vegetation types. Coordinated and standardized efforts, such as the NEON network, will help to overcome part but not all of these issues (Kamoske et al., 2022).

Pacheco-Labrador et al. (2022) demonstrated that FDMs with low performance in global-scale comparisons yielded a wide range of results at local scales. They also showed that uncertainty spuriously inflated the performance of certain metrics, and how specific preprocessing (standardization and Principal Component Analyses) was necessary to establish relationships between plant traits and remote sensing-derived FDMs. Simulations provide a theoretical framework for testing methodological hypotheses and benchmarking metrics and methods (Fassnacht et al., 2022; Ludwig et al., 2024) before they are applied to real-world observations. This potential led to the creation of the first Biodiversity Observing System Simulation Experiment (BOSSE v1.0), designed to support the study of plant functional diversity and biodiversity-ecosystem function relationships (https://github.com/JavierPachecoLabrador/pyBOSSE) (Pacheco-Labrador et al. 2025). BOSSE generates synthetic scenes that include all plant traits (PT) and optical spectral signals such as hyperspectral reflectance factors (*R*), derived optical traits (OT), sun-induced chlorophyll fluorescence radiance (*F*), and land surface temperature (LST), linked via radiative transfer theory, necessary to test new methodologies. The model is open source and available to the scientific community to support advances in remote sensing of plant functional diversity.

This manuscript presents the first use of BOSSE to test five fundamental, yet unsolved methodological questions, each with an underlying hypothesis about the outcome. We compare the simulation results with previous findings and offer recommendations and guidance for estimating plant functional diversity from remote sensing.

## 2. METHODOLOGICAL QUESTIONS AND HYPOTHESES

### Q1. How should remote sensing characterize functional diversity in large surfaces (sites)?

Many works have computed FDMs over relatively small patches or moving windows (Cimoli et al., 2024; Tagliabue et al., 2020), in some cases matched with field plots for validation (Hauser et al., 2021). However, it is not yet clear how to use remote sensing to characterize the diversity of large surfaces, which could be a study “site” (or several), understood as an area sufficiently large to evaluate the role of biodiversity on ecosystem functions. The size of the area where diversity is computed presents a saturating relationship with diversity (Ricotta et al., 2012; Walker et al., 2008); this rarefaction effect also occurs in remote sensing (Gholizadeh et al., 2022). However, while the impact of the window size used to compute the FDMs has been reported (Cimoli et al., 2024; Schneider et al., 2017), the effect on the comparison of field and remote sensing FDMs remains unexplored. Thus, at the site scale, FDMs could be computed to the whole extent of the study area, or as a central trend of smaller plots (e.g., moving windows) within such an area.

H1: We hypothesize that averaging the diversity of small windows might better connect plant traits and remote sensing-based FDMs.

### Q2. Which sources of spectral information (or combinations thereof) of spectral variables can best capture plant functional diversity at the site scale?

There is an intrinsic variability in the relationships between the signals captured by remote sensors and plant properties due to the differences in the source (reflection or emission) and the wavelength-dependent nature of light-matter interactions governing the different spectral signals (e.g., Verrelst et al. (2015)). Since plant traits correlate differently with each spectral signal or variable, some of these might be better estimators of the diversity of the plant traits. While former local works compared different spectral regions or spectral resolutions (Wang et al., 2018a) or spectral indices and sun-induced chlorophyll fluorescence (Tagliabue et al., 2020), a systematic comparison of all passive optical remote sensing signals and optical traits at a broad range of situations has not yet been performed.

H2: We hypothesize that some of the variables that passive optical remote sensing can provide, alone or in combination, can better capture plant functional diversity than others and should be preferentially used.

### Q3. How should remote sensing estimates be validated/compared with plant functional diversity measurements?

Pixel area has been proposed as a proxy to represent species’ relative abundance when computing FDMs from remote sensing imagery (Laliberté et al., 2020; Rocchini et al., 2017; Rossi et al., 2021b). However, given the control leaf area index (LAI) has on the radiative transfer, it could be a better proxy of the species (or pixel) abundance when validating or comparing remote sensing and field plant trait diversity estimates.

H3: We hypothesize that if used as an abundance surrogate, LAI should be able to provide FDM values that are more comparable between field and RS-based estimates.

### Q4. When (in the phenological year) can remote sensing best capture site-scale plant functional diversity?

Compared to fieldwork sampling, the recurrent nature of remote sensing enhances possibilities for assessing diversity in the temporal domain (Rossi et al., 2021b). However, there is likely a better time to catch the underlying diversity since confounding factors might hide the variability of vegetation traits at specific phenological periods, as background contribution (Wang et al., 2018b) or the saturation of some spectral signals produced by vigorous vegetation (Gao et al., 2023). It thus remains unclear how plant phenology affects the uncertainty in estimating plant functional diversity.

H4: We hypothesize that intermediate states of vegetation development (during development or decay) might be optimal for characterizing plant functional diversity.

### Q5. Which approaches and remote sensing variables are more resistant to the effects of suboptimal spatial resolution?

Spatial resolution poses substantial limitations to the estimation of plant diversity from remote sensing, as reported in multiple studies (Helfenstein et al., 2022; Pacheco-Labrador et al., 2022; Torresani et al., 2024; Wang et al., 2018b). Pacheco-Labrador et al. (2022) showed that correlations are stronger when, under suboptimal spatial resolution, FDMs were calculated from field plant traits aggregated to the same resolution of the remote sensor (“sameSR”) than per individual or species basis (“highSR”). However, broader assessments are lacking, and it remains unclear whether specific methodologies, remote sensing variables, or approaches to compare remote sensing and field plant trait diversity estimates are more robust to the degradation of spatial resolution. Based on previous findings, the resilience to suboptimal spatial resolution depends on the remote sensing variable used to estimate the FDM and the proxy of abundance used to calculate FDMs from the field plant traits (“per-pixel” or “per-LAI”).

H5.1: We hypothesize that the best-performing remote sensing variables (see Q2) will also be the most resilient to the effects of suboptimal spatial resolution.

H5.2: We hypothesize that “per-LAI” would provide the best performance for suboptimal (< 100 %) spatial resolutions.

H5.3: We hypothesize that degrading field plant trait data to the imagery resolution is necessary to estimate plant functional diversity from space remote sensing.

## 3. MATERIAL AND METHODS

### 3.1 BOSSE simulations

#### 3.1.1. BOSSE Scenes

BOSSE generates “Scenes” encompassing individuals of different species (Fig. 1b,g,l,q) assigned to different plant functional types (Fig. 1a,f,k,p). Each pixel of the Scene represents an individual or a group of identical individuals of the same species, whose plant traits evolve in response to meteorology. Therefore, BOSSE pixels do not have an absolute size and are instead defined as units of homogeneous vegetation, and the spatial resolution is described in relative terms (pixel to plant size). BOSSE species are defined by: 1) a set of upper and lower bound ranges of plant functional traits (i.e., biophysical and physiological variables of the model SCOPE (van der Tol et al., 2009)); and 2) a set of coefficients of the growing season index model developed by Forkel et al. (Forkel et al., 2014) that determine their growth sensitivity to different environmental factors (i.e., air temperature, soil water availability, and incoming radiation). The species exhibit intra-specific variability in trait bounds, which are overall delimited by plant function, resulting in each pixel having unique values (Fig. 1c, h, m, r). The trait values of each species’ individual at any point in time are scaled within their bounds using the growing season index (which ranges between 0 and 1) at that time (Fig. 1d,l,n,s). BOSSE utilizes emulators of the SCOPE model to enable fast simulation of remote sensing imagery from plant trait maps and meteorological data (Fig. 1e,j,o,t).

**Figure 1.**
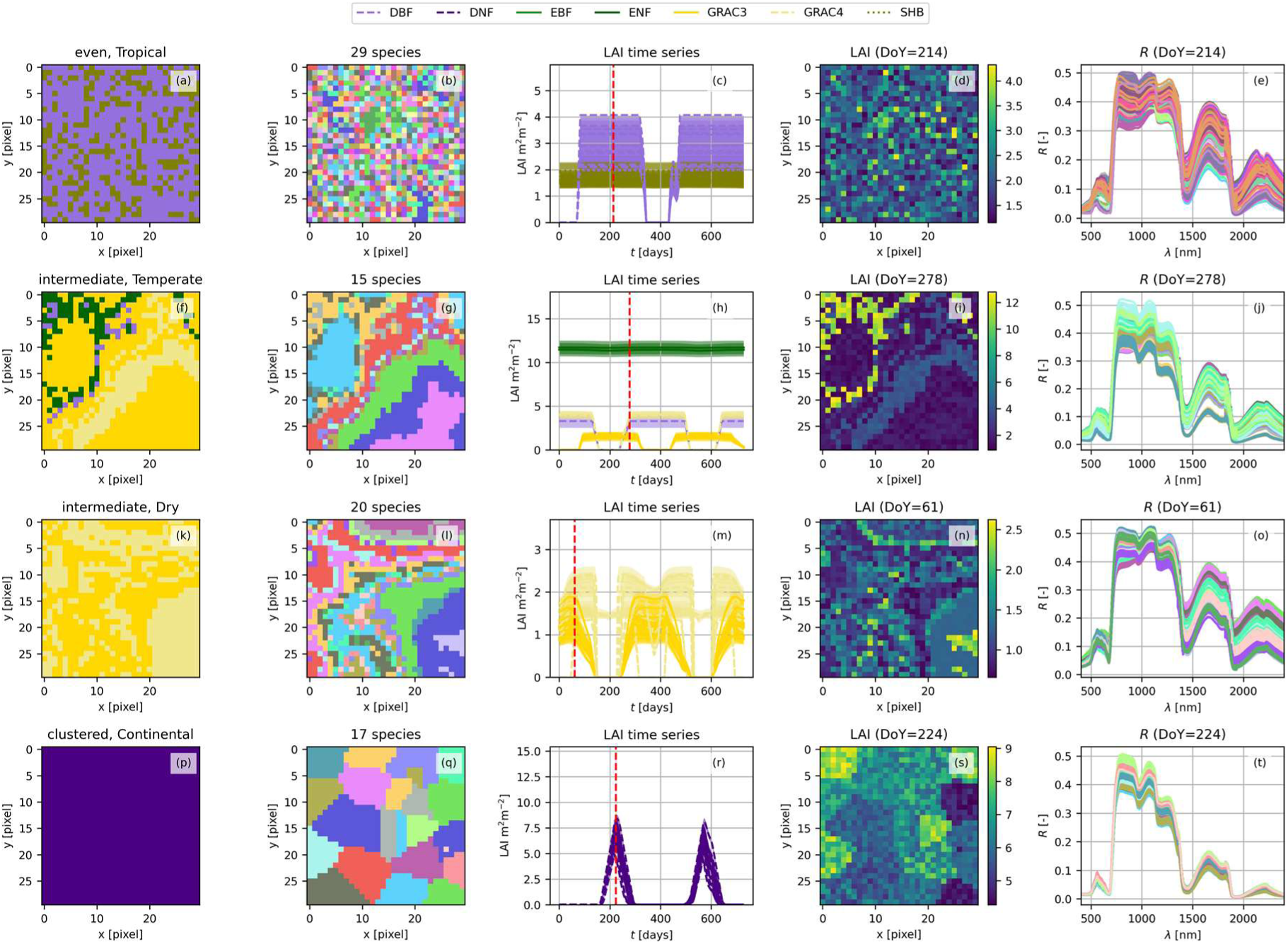
Examples of four BOSSE Scenes simulated for the different climatic zones: Tropical (a-e), Temperate (f-j), Dry (k-o), and Continental (p-t), and spatial patterns: even (a-e), intermediate (f-o), clustered (p-t). For each Scene, the figure presents the plant functional types map (first column, see legend for color code), the species map (each species represented by a different color), the time series of leaf area index (mean and standard deviation interval of each species, colored by plant functional type (third column, see legend for color code), the leaf area index map at the date of the maximum phenological development (fourth column), and the hyperspectral reflectance factor of each pixel, coloured by species.

Table 1 summarizes the plant traits simulated by BOSSE and used to compute FDMs (see section 2.1.2) and/or simulate spectral imagery. See Pacheco-Labrador et al. (2025) for a detailed description of the BOSSE design and simulation approach.

**Table 1.**
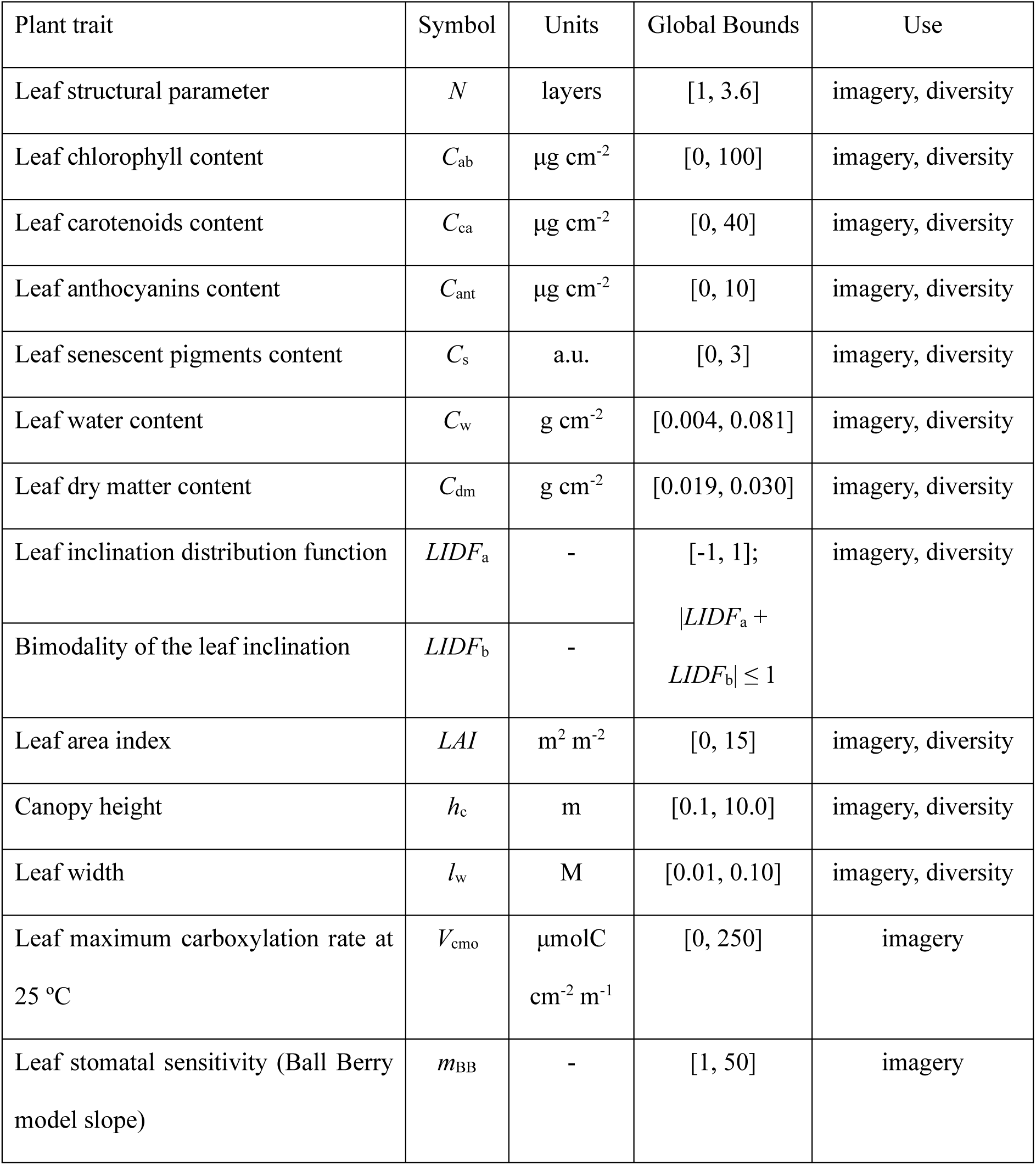
Plant traits simulated by BOSSE and used for simulating remote sensing imagery (“imagery”) or calculating diversity (“diversity”).

We simulated 15 “Scenes” under different meteorological conditions in four different climatic zones (“Tropical”, “Dry”, “Temperate”, and “Continental”) with three other species spatial pattern distributions (“clustered”, “intermediate”, and “even”) (e.g., Figure 1). In total, 180 Scenes. BOSSE takes as input meteorological data corresponding to a 2-year-long ERA-5 hourly time series available in Zenodo (Pacheco-Labrador et al. 2025). For each Scene, we generated plant trait maps and remote sensing imagery every four days at midday (a frequency similar to that of Sentinel-2), from which we computed FDMs. Scenes are 30-by-30 pixels, assuming that each pixel corresponds to one individual or a group of identical individuals from the same species. Thus, simulations are initially performed at 100% relative spatial resolution; however, this resolution can then be degraded for plant trait maps and remote sensing imagery for analysis, where pixels combine signals from individuals of different species.

In addition, we assessed the effect of the window size used to compute the FDMs on a different dataset that encompassed 180 Scenes of 90-by-90 pixels for the same meteorological time series, climatic zones, and spatial patterns. In this case, we generated trait maps and imagery only at the peak of each year’s season (i.e., maximum averaged growing season index), at midday.

#### 3.1.2. Functional diversity metrics

Based on a previous assessment of FDMs linking plant trait-based and remote sensing-based diversity (Pacheco-Labrador et al., 2022), we computed Rao’s quadratic entropy (*Q*_Rao_) to represent *α*-diversity, and used the variance-based partitioning method (Laliberté et al., 2020) to estimate diversity at different scales *α* and *β* (i.e., within and between the moving windows, respectively). By default, 3-by-3 windows were used to compute these metrics. To enable a direct (1:1) comparison of the FDMs computed from the various plant trait and remote sensing datasets, each featuring a different number of variables, we used the “pyGNDiv” package (https://github.com/JavierPachecoLabrador/pyGNDiv-master) (Pacheco-Labrador et al., 2023). This package normalizes the dissimilarity metric (Euclidean distance for *Q*_Rao_ and the sum of squares for the variance-based partitioning) to yield dimensionality-independent metrics. Unlike previous simulation works (Pacheco-Labrador et al., 2023, 2022), BOSSE explicitly describes the spatial distributions of species and remote sensing pixels. Therefore, FDMs were computed within squared moving windows that scanned the Scene maps and imagery as proposed by Rocchini et al. (Rocchini et al., 2017). To achieve this, we utilized an updated version of the “pyGNDiv” package, which includes parallelization and image processing capabilities, available on GitHub.

The FDMs were computed for plant trait maps (PT), representing the ideal case for field data where all individual plant traits have been sampled in the Scene, and for different combinations of remote sensing products (Table 2). We only considered as PT those biophysical variables that had a physical direct interaction with radiation (thus, excluding *V*_cmo_ and *m*_BB_, Table 1). The same variables were estimated from the reflectance imagery as the remote sensing-based optical traits (OT). Both for PT and OT, we did not calculate FDMs in any moving window that included pixels without vegetation (LAI = 0.0 for PT, LAI ≤ 0.1 for OT). The vegetation’s presence/absence information would be available from the simulation (equivalent to field measurements) or estimated from the OT retrieval. Furthermore, we assumed no vegetation when *F* ≤ 0.1 mW m^-2^ sr^-1^ nm^-1^. We set the LAI and *F* threshold values larger than zero to accommodate some retrieval uncertainty. However, this prior knowledge was not assumed for the other spectral signals (*R* and LST), except when combined with OT and/or *F*, and bare-soil pixels were included in the computation. Spectral indices can summarize a large fraction of the information contained in *R*. Thus, for comparison, we computed two spectral indices related to vegetation density and greenness, the Normalized Difference Vegetation Index (NDVI) (Kriegler et al., 1969), commonly used, and the Near Infrared of Vegetation (NIR_v_) (Badgley et al., 2017), designed to be more robust to background effects than NDVI. Moreover, for plant and optical traits, we tested using LAI (“per-LAI”) or the ground area (in pixels, “per-pixel”) of each pixel as a proxy of species abundance. In the first case, LAI was not used as a plant functional trait to compute the FDMs. When combining different spectral signals, we assumed they featured the same spatial resolution, which is not realistic since *F* and LST satellite imagery usually present lower resolutions than multispectral or hyperspectral *R*. This is because we wanted to test the potential capability of each signal to inform about plant functional diversity. The effect of the differences in the spatial resolution is discussed (section 4) and can be assessed with BOSSE in future studies.

**Table 2.**
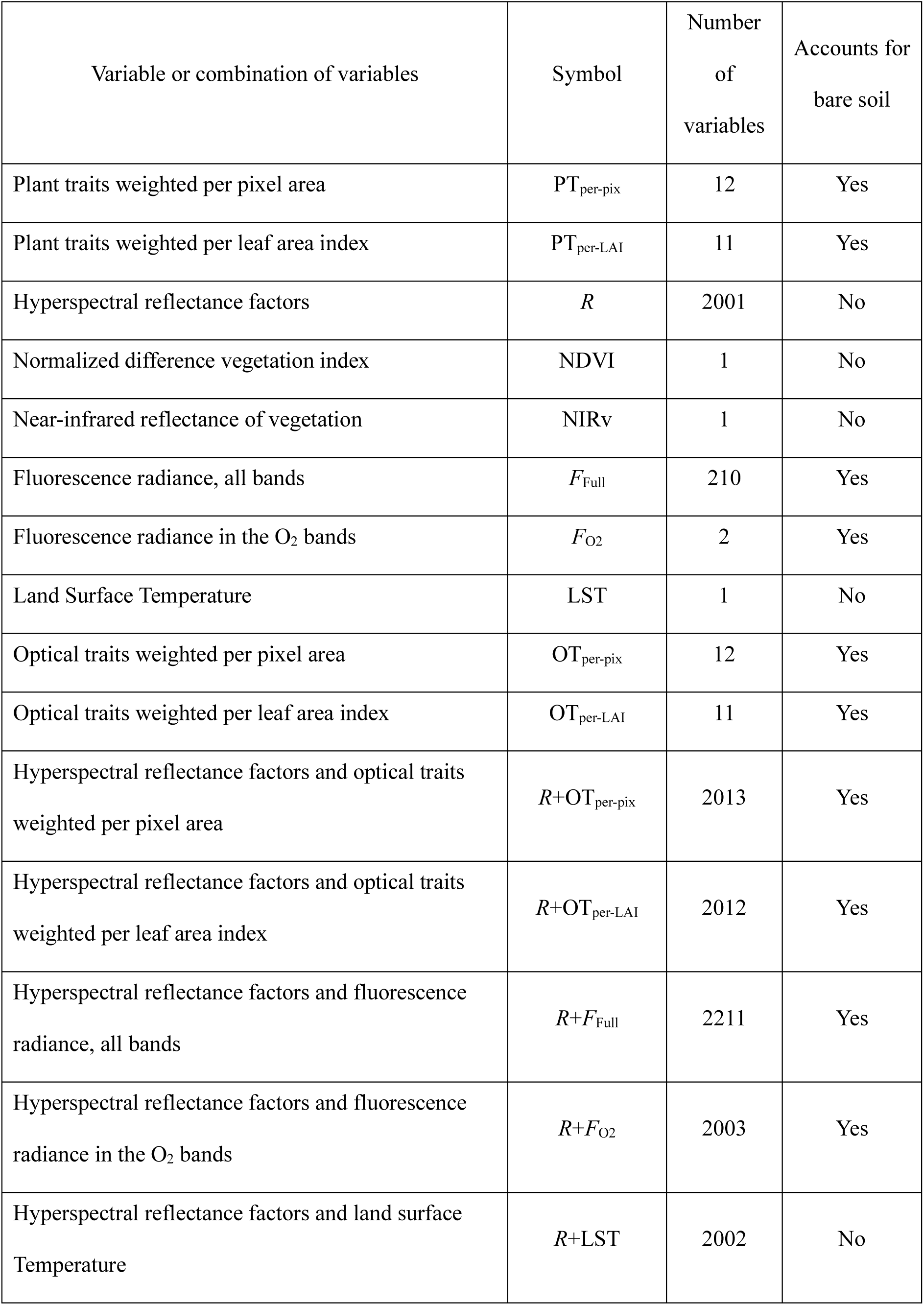
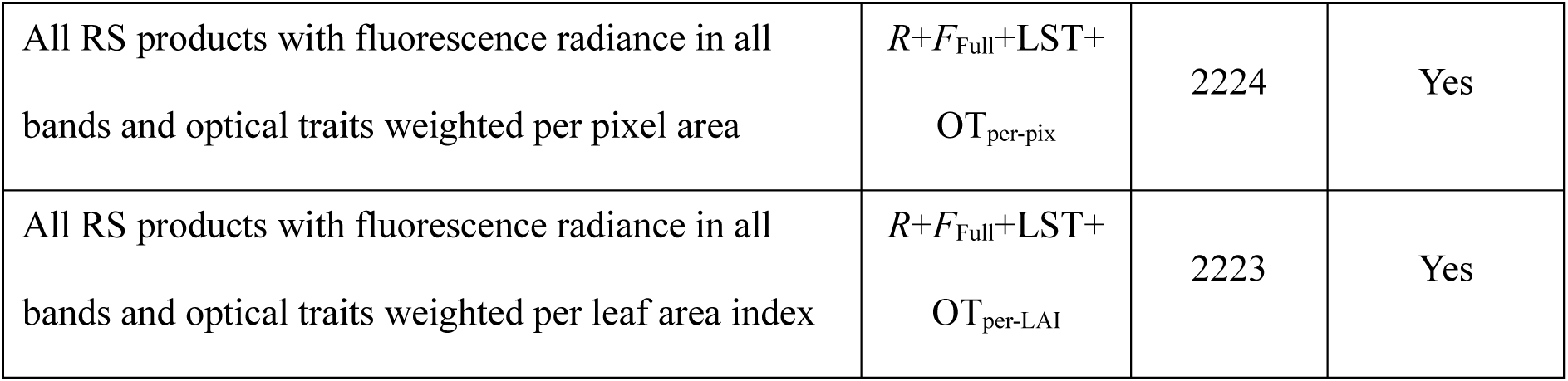
Plant traits, remote sensing variables, or combinations of variables used to compute functional diversity metrics.

| Variable or combination of variables | Symbol | Number of variables | Accounts for bare soil |
| --- | --- | --- | --- |
| Plant traits weighted per pixel area | $PT_{\text{per-pix}}$ | 12 | Yes |
| Plant traits weighted per leaf area index | $PT_{\text{per-LAI}}$ | 11 | Yes |
| Hyperspectral reflectance factors | $R$ | 2001 | No |
| Normalized difference vegetation index | NDVI | 1 | No |
| Near-infrared reflectance of vegetation | NIR<sub>v</sub> | 1 | No |
| Fluorescence radiance, all bands | $F_{\text{Full}}$ | 210 | Yes |
| Fluorescence radiance in the O<sub>2</sub> bands | $F_{\text{O}_2}$ | 2 | Yes |
| Land Surface Temperature | LST | 1 | No |
| Optical traits weighted per pixel area | $OT_{\text{per-pix}}$ | 12 | Yes |
| Optical traits weighted per leaf area index | $OT_{\text{per-LAI}}$ | 11 | Yes |
| Hyperspectral reflectance factors and optical traits weighted per pixel area | $R+OT_{\text{per-pix}}$ | 2013 | Yes |
| Hyperspectral reflectance factors and optical traits weighted per leaf area index | $R+OT_{\text{per-LAI}}$ | 2012 | Yes |
| Hyperspectral reflectance factors and fluorescence radiance, all bands | $R+F_{\text{Full}}$ | 2211 | Yes |
| Hyperspectral reflectance factors and fluorescence radiance in the O<sub>2</sub> bands | $R+F_{\text{O}_2}$ | 2003 | Yes |
| Hyperspectral reflectance factors and land surface Temperature | $R+LST$ | 2002 | No |
| All RS products with fluorescence radiance in all bands and optical traits weighted per pixel area | $R+F_{\text{Full}}+\text{LST}+$<br>$\text{OT}_{\text{per-pix}}$ | 2224 | Yes |
| All RS products with fluorescence radiance in all bands and optical traits weighted per leaf area index | $R+F_{\text{Full}}+\text{LST}+$<br>$\text{OT}_{\text{per-LAI}}$ | 2223 | Yes |

### 3.2 Addressing the research questions

Q1. How should remote sensing characterize functional diversity in large surfaces (sites)?

We address the issue of estimating diversity in a “site”, that is, a large extent where biodiversity could be relevant for ecological processes. The sites are represented in this case by the simulated Scenes, 30 pixels long. We compare field and RS-estimated FDMs computed over a unique window of 30-by-30 pixels (“scene” approach) and the mean or the median of an FDM computed over 3-by-3 pixel moving windows (“mean” or “median” approaches). The comparison is performed “between sites” (by comparing the temporally-averaged FDMs of the 180 Scenes simulated) and “within sites” (by comparing the FDMs computed at each time stamp of the simulation, site by site). In addition, we assessed the effect of window size on a different dataset of larger Scenes (90-by-90 pixels) focused on each year’s growing season peak, and computed FDMs using moving windows of 3, 5, 10, 15, 30, 45, and 90 pixels (the entire Scene).

Q2. Which sources of spectral information (or combinations thereof) of spectral variables can best capture plant functional diversity at the site scale?

We test the capability of different remote sensing variables and combinations of *R* with OT, *F*, and/or LST to capture plant functional diversity as computed from the simulated vegetation traits (Table 2). FDMs from field plant traits and remote sensing-estimated optical traits are calculated using pixel coverage or LAI as a surrogate of abundance. However, when comparing them, the terms “per-pixel” or “per-LAI” refer to the method by which the field plant traits-based FDMs are computed.

Q3. How should remote sensing estimates be validated/compared with plant functional diversity measurements?

We compare using LAI as a surrogate of the species’ abundance (“per-LAI”) with the case where pixel area is the proxy for abundance and LAI is used as a functional trait (“per-pixel”) to link field- and remote sensing-estimated FDMs (listed in Table 1). Both approaches are applied to field plant traits and to the optical traits, where LAI is also retrieved from *R*.

Q4. When (in the phenological year) can remote sensing best capture site-scale plant functional diversity?

We assess the dependency of the FDM estimation error on the phenological status, represented by LAI (which is driven by the growing season index), at each timestamp of all the simulations.

Q5. Which approaches and remote sensing variables are more resistant to the effects of suboptimal spatial resolution?

We compare field plant trait- and remote sensing-estimated FDMs at different spatial resolutions, defined in relative terms as the ratio of the plant to the pixel size (100%, 90%, 60%, 30%, and 10%). Furthermore, we compare the remote sensing estimates with vegetation diversity calculated from traits aggregated at the suboptimal remote sensor resolution (“same SR”) and at the highest resolution (“high SR”). We also compare the remote sensing estimates with field estimates weighted “per-LAI” and “per-pixel”.

### 2.3 Statistical analyses

Statistical analyses were performed in Python 3.11. We compared the FDMs computed from plant trait maps and the different remote sensing signals (Tables 1 and 2) using the coefficient of determination (*R*^2^), to capture the accuracy and precision of the estimation, the Pearson correlation coefficient (*r*^2^), to assess the strength of the relationship independently of biases, and the root mean squared error (RMSE) to evaluate the difference between metrics (Katja Richter et al., 2012). For each pair of field plant trait- and remote sensing-estimated FMD, we computed the statistics comparing: 1) the mean FDM of each synthetic site (averaged for the time series, “between sites”) for all the Scenes; and 2) the FDM of each date for each individual Scene (“within sites”).

## 3. RESULTS

### Q1. How should remote sensing characterize functional diversity in large surfaces (sites)?

Out of the three approaches explored (“mean”, “median”, or “scene”), the “mean” and “median” of small moving windows outperformed the “scene” approach in estimating plant functional diversity (i.e., *Q*_Rao_) at the site scale. This result appears to be due to the saturating relationship between the diversity metrics and the number of pixels used to compute them, which reduces the correlation between field plant traits and remote sensing estimates of diversity. Consequently, we exclude the “scene” approach from the subsequent analyses since it offers no advantage (not shown).

Fig. 2 presents the distributions of the statistics computed for all the remote sensing metrics “between sites” (i.e., comparing the *Q*_Rao_ temporal mean value of each site). “Mean” and “median” show higher performance and lower variability between the remote sensing predictors than the “scene” approach. However, the performance decreases and the variability increases when *Q*_Rao_ time series are compared individually at each site (“within sites”, Fig. S1). These results suggest that the capability of remote sensing to capture plant functional diversity at a given site can strongly depend on the site’s characteristics.

**Figure 2.**
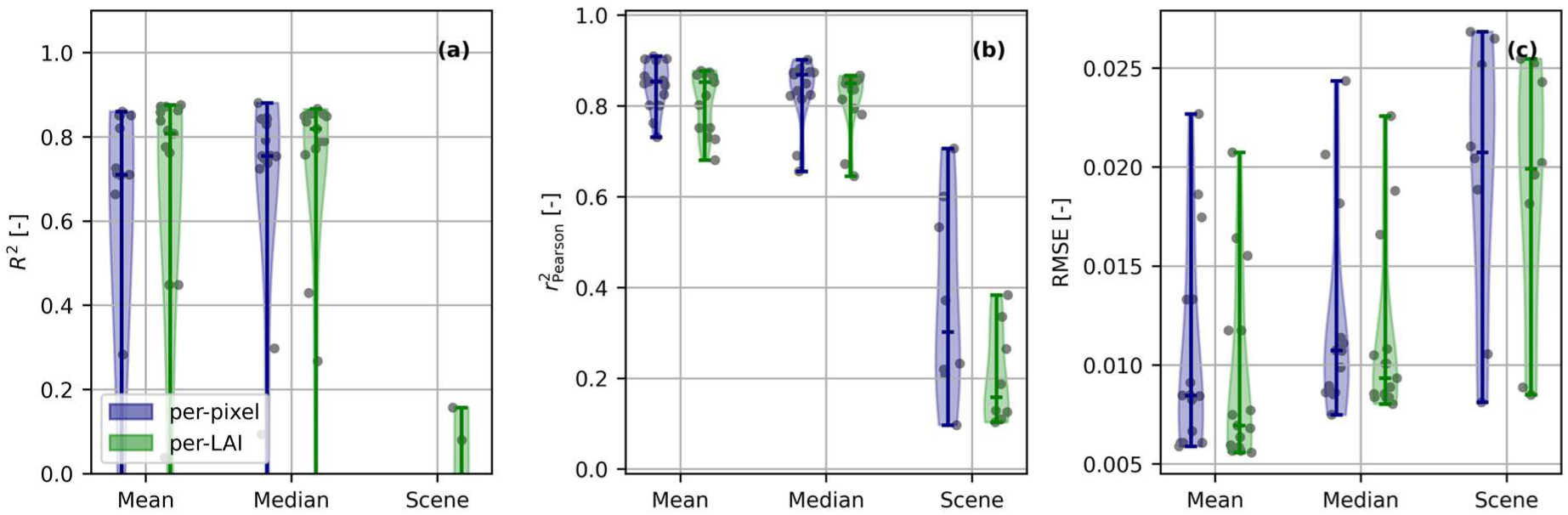
Distribution of the statistics resulting from the comparison of functional diversity metrics computed from plant traits and several combinations of remote sensing variables. Plant traits’ abundance is represented by different proxies: pixels covered (“per-pixel”) or leaf area index (“per-LAI”). Coefficient of determination (a), Pearson correlation coefficient (b), and root mean squared error (c). The comparison is performed “between sites” by comparing the temporally-averaged Rao’s quadratic entropy index (*Q*).

Small moving windows (i.e., 3-by-3 pixels) maximize the differences of *Q*_Rao_ and *f*_α_ between spatial patterns (Fig. 3), which present increasing values from clustered (Fig. 1p-t) to even (Fig. 1a-e). However, their values reach a saturation point at window sizes between 30-by-30 and 45-by-45 pixels, converging to similar values regardless of the spatial distribution of species. This process minimizes the range of variability of the metrics and the correlation strength between field plant traits and remote sensing estimates (Fig. 3, S2 and S3). For diversity partitioning, *f*_α_ reaches 100% once the window size matches the Scene size.

**Figure 3.**
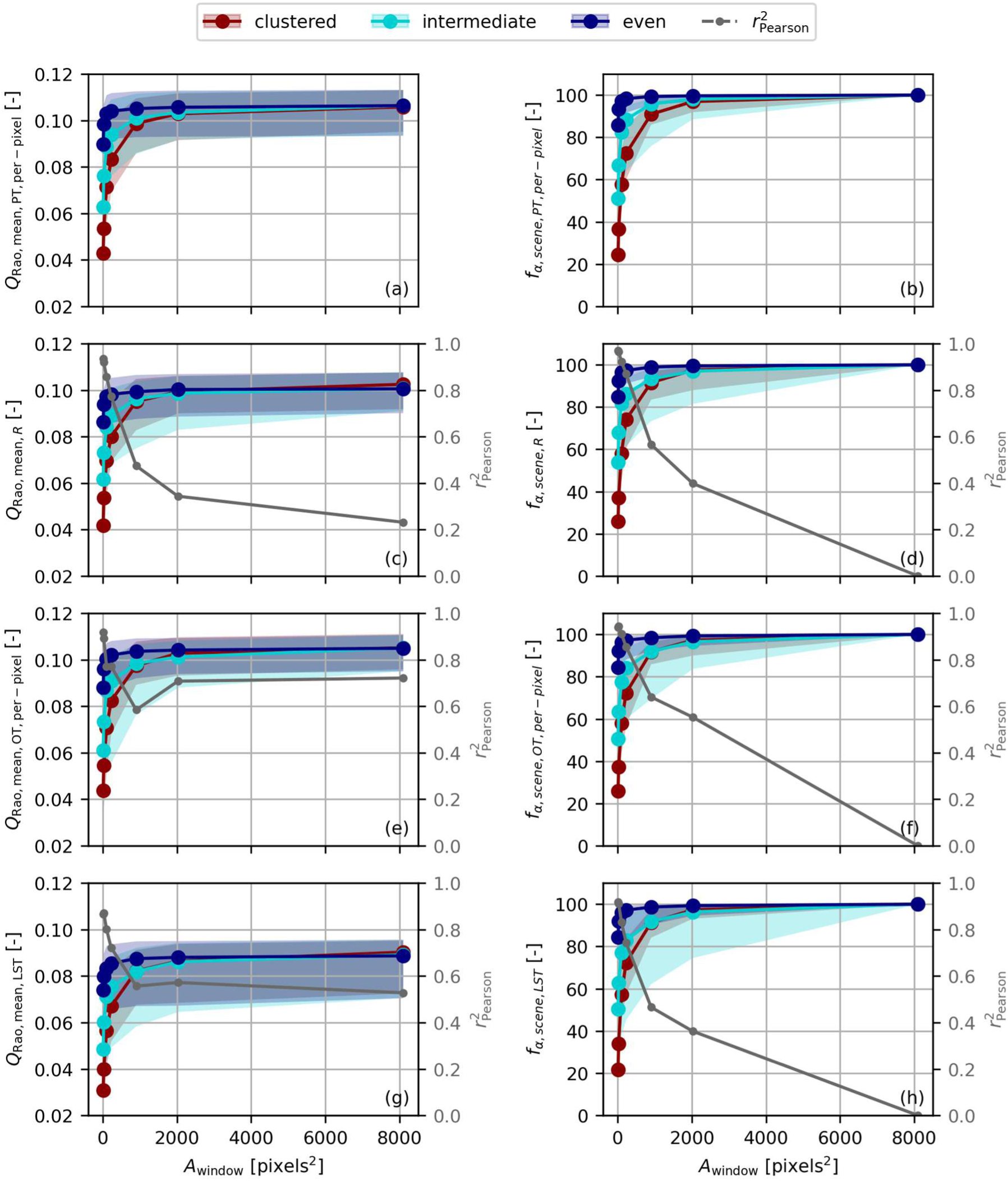
Functional diversity vs. moving window area (*A*_window_) curves separated by the spatial pattern of vegetation distribution type. Functional diversity is computed as the mean of the *Q*_Rao_ of small moving windows.

### Q2. Which sources of spectral information (or combinations thereof) of spectral variables can best capture plant functional diversity at the site scale?

OT and *F*, alone or in combination with other variables, result in the best estimators of plant functional diversity (*Q*_Rao_, Fig. 4a-f), while LST yields the worst results. Compared to the “mean” approach, the “median” approach improves the correlation (*R*^2^ and *r*^2^*_pearson_*) of the least performing variables, but increases RMSE except for *R* and *R*+LST (Fig. 4c). The results are consistent for variance-based diversity partitioning (Fig. 4d-f), where the least performing variables are the same as for *Q*_Rao_. The correlation statistics (*R*^2^ and *r*^2^*_pearson_*) are much lower and present a wide variability when individual sites (“within sites”, Fig. S4), but RMSE values are similar for *Q*_Rao_ (Fig. S4c,f,i,l), and lower for variance-based diversity partition (Fig. S4o,r).

**Figure 4.**
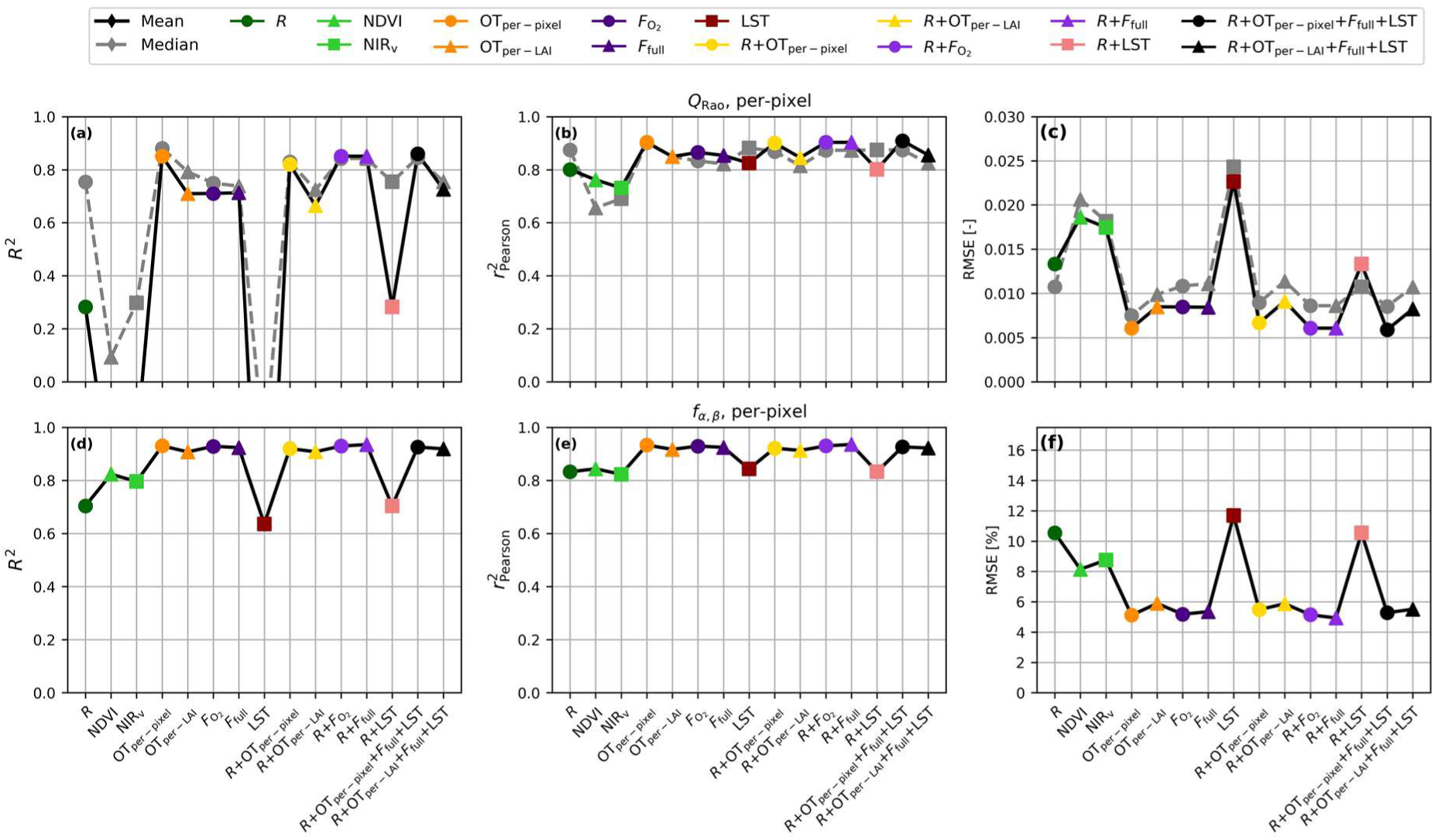
Statistics resulting from the comparison of functional diversity metrics computed from plant traits and several combinations of remote sensing variables using the mean of small moving windows and using pixel area as a proxy of abundance: for *Q*_Rao_ (a-c), and for a fraction of alpha and beta-diversity (d-f). Coefficient of determination (a, d), Pearson correlation coefficient (b, e), and root mean squared error (c, f).

### Q3. How should remote sensing estimates be validated/compared with plant functional diversity measurements?

Comparing remote sensing-estimated *Q*_Rao_ with field plant trait estimates weighted by LAI (“per-LAI”) increases the accuracy and precision of the estimation, but reduces the correlation strength (*r*^2^*_pearson_*) for most remote sensing variables. However, for variance-based diversity partitioning (*f_α_*_,*β*_), “per-LAI” abundance leads to varying results, often reducing correlation and increasing error.

Fig. 5 presents the difference between the statistics of both approaches (“per-LAI” minus “per-pixel”). For *Q*_Rao_, “per-LAI” abundance increases *R*^2^ (Fig. 5a) and reduces RMSE (Fig. 5c), particularly for the least performing variables (LST, NDVI, and NIR_v_, Fig. 4). Nonetheless, this increase in accuracy occurs at the expense of a reduction in the correlation strength (*r*²_Pearson_, Fig. 5b) for all variables, except for *Q*_Rao_ computed from OT weighted “per-LAI”, alone or combined with other variables (Fig. 5b). Regarding variance-based diversity partitioning (*f_α_*_,*β*_) “per-LAI” abundance reduces *R*^2^ (Fig. 5a), *r*^2^ (Fig. 5b), and increases RMSE (Fig. 5c) except for some of the least performing variables (*R*, LST, and *R*+LST). The differences are similar when comparing the statistics computed site by site (“within sites”, Fig. S5). In most cases, *R*^2^ increases (Fig. S5a) but *r*^2^*_pearson_* decreases (Fig. S5b); RMSE increases for *Q*_Rao_ and decreases for variance-based diversity partitioning (Fig. S5c).

**Figure 5.**
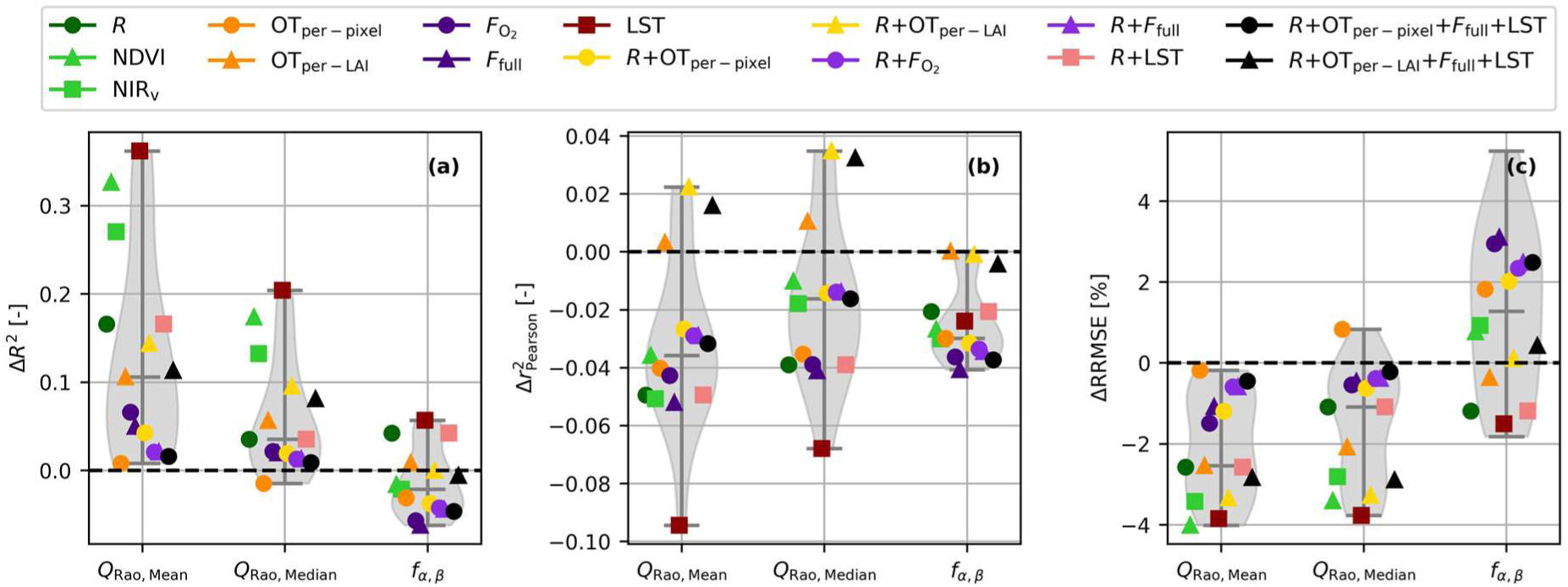
Difference between the statistics resulting from the comparison of remote sensing-based and field-based functional diversity metrics. The statistics correspond to the comparison of the mean metric values across all sites. The difference (*Δ*) is built as the difference of the statistics for the case where field-based abundance is represented by the leaf area index (“per-LAI”) minus the case where it is represented by the pixel coverage (“per-pixel”). Coefficient of determination (a), Pearson correlation coefficient (b), and root mean squared error (c). The statistics represent a comparison of the temporally averaged diversity metrics of all the simulated scenes (“between sites”).

### Q4. When (in the phenological year) can remote sensing best capture site-scale plant functional diversity?

The estimation of plant functional diversity achieves low and stable RMSE once vegetation develops (i.e., LAI > 1), particularly when bare pixels are not identified and removed from remote sensing data. The “per-LAI” approach often minimizes RMSE at the time of maximum development (i.e., LAI).

Fig. 6 summarizes the mean Scene LAI–RMSE relationship for the “per-LAI” approach (see Fig. S6 for the “per-pixel” approach); each data point corresponds to a Scene and date, the lines correspond to the smoothed curves, and red stars locate the minimum RMSE of each curve. Field *Q*_Rao_ (Fig. 6a-h, S6a-h) and *f_α_* or *f_β_* (Fig. 6i-o, S6i-o) RMSE asymptotically decreases with the averaged Scene LAI, being RMSE maximum at low LAI (e.g, LAI < 1). When field data uses “per-LAI” abundance, Q_Rao_ RMSE minimizes at the maximum LAI for *R*, NIR_v_, OT_per-LAI_ alone and combined with other variables (Fig. 6a-e), and falls between values of 3 and 4 for the rest. The “per-pixel” abundance minimizes the *Q*_Rao_ RMSE with a maximum LAI for NIR_v_ and OT, falling the rest of the RMSE minima between LAI values of 1 and 3.5 (Fig. S4a-e). For variance-based diversity partitioning, RMSE minimizes at maximum LAI for NIR_v_, OT, and *F* for both abundance approaches, and also for R+*F*_full_ for the “per-LAI” approach (Fig. 6i-p, S4i-p), with the rest of the RMSE minima falling between LAI values of 3 and 4. Nonetheless, most RMSE-mean LAI relationships flatten for LAI > 1 or even lower, making the RMSE differences across this range small.

**Figure 6.**
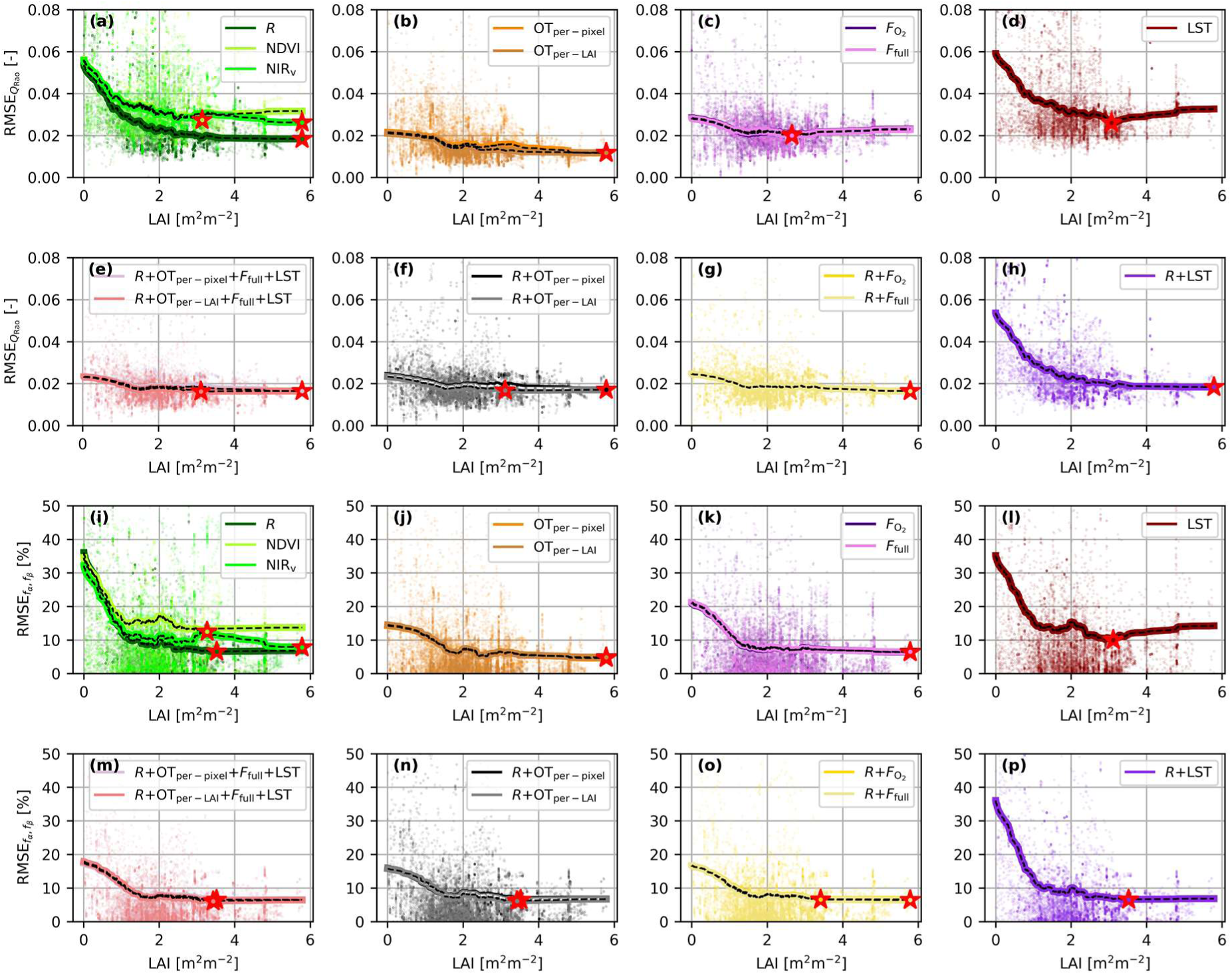
Relationship between mean scene leaf area index (LAI) and the root-mean-squared error of the plant functional diversity (Rao’s quadratic entropy index (*Q*_Rao_)) and the fractions of alpha and beta-diversity (combined) at each simulation timestamp. Field-based functional diversity is computed using LAI for abundance (“per-LAI”), and it is estimated with different remote sensing sets of variables: reflectance factors (*R*), optical traits used to calculate diversity using the pixel area (*R*+OT_per-pix_) or leaf area index (*R*+OT_per-LAI_) for abundance, sun-induced chlorophyll fluorescence radiance retrieved at the O_2_ absorption bands (*F*_02_), the entire emission spectrum (*F*_full_), land surface temperature (LST), and combinations of the former ones.

### Q5. Which approaches and remote sensing variables are more resistant to the effects of suboptimal spatial resolution?

The estimation of *Q*_Rao_ from remote sensing is severely affected by spatial resolution, particularly when remote sensing estimates are compared with field estimates at high spatial resolution (“high SR”). Matching remote and field data resolutions (“same SR”) reduces sensitivity and improves correlations. The best estimators (see Q2) are also the most robust. For variance-based partitioning, there are smaller differences between “high SR” and “same SR”, resulting in better performance than *Q*_Rao_. In all cases, the estimation is unreliable for relative spatial resolutions smaller than 30% (i.e., pixel sizes three times larger than the plant size).

Fig. 7 summarizes the sensitivity of the different spectral variables to spatial resolution when using LAI as a proxy of field data abundance (“per-LAI”, see Fig. S7 for the “per-pixel” approach). Fig. 7 separates variables that include (columns 3-4) or exclude (columns 1-2) hyperspectral reflectance factors, as well as the “same SR” (columns 1-3) and “high SR” (columns 2-4) comparisons. For *Q*_Rao_ (Fig. 7a-l), the “high SR” *r*^2^*_pearson_* decreases as spatial resolution coarsens, whereas for “same SR”, *r*^2^*_pearson_* slightly increases at mild suboptimal resolutions (90-60 %). The impact of spatial resolution on the estimation accuracy is mixed, particularly for *R*^2^ (Fig. 7a-d). *R*^2^ slightly increases at mild suboptimal resolutions for “same SR”, while it decreases and then recovers around 60% for “high SR”. The effect on RMSE (Fig. 7i-l) is more consistent between approaches (“same SR” or “high SR”); in all cases, “same SR” RMSE minimizes at the lowest resolution, whereas “high SR” RMSE maximizes (Fig. 7k,l). *R*^2^ is less sensitive to substantial degradation of <u>spatial</u> resolution (i.e., less than 60 %) for the “per-LAI” approach (Fig. 7a-d vs. Fig. S7a-d), whereas it is more similar for *r*^2^ and RMSE. Among the “No R-based” variables (Fig. 7a-b,e-f,i-j), OT and *F* are more robust to suboptimal spatial resolution than LST and the spectral indices in all cases. Regarding “*R*-based” variables (Fig. 7c-d,g-h,k-l), *R* alone features an intermediate performance, which improves when combined with OT and/or *F*. Overall, when the pixel size is more than three times larger than the plant size, none of the remote sensing signals and approaches tested can capture plant trait *Q*_Rao_ (Fig. 7a-l). The differences between remote sensing signals and methods are less significant for variance-based diversity partitioning (Fig. 7m-x, Fig. S7m-x). In this case, “same SR” does not necessarily perform better than “high SR”; in fact, “high SR” achieves the largest accuracies (*R*^2^, Fig. 7m-p), lowest errors (RMSE, Fig. 7u-v), and strongest correlations (*r*^2^*_pearson_*, Fig. 7q-t) for the “per-LAI” approach. Alternatively, “same SR” performs best for the “per-pixel” approach (Fig. S7m-p). Regarding the variables used, *R*, LST, and *R*+LST show the lowest performances.

**Figure 7.**
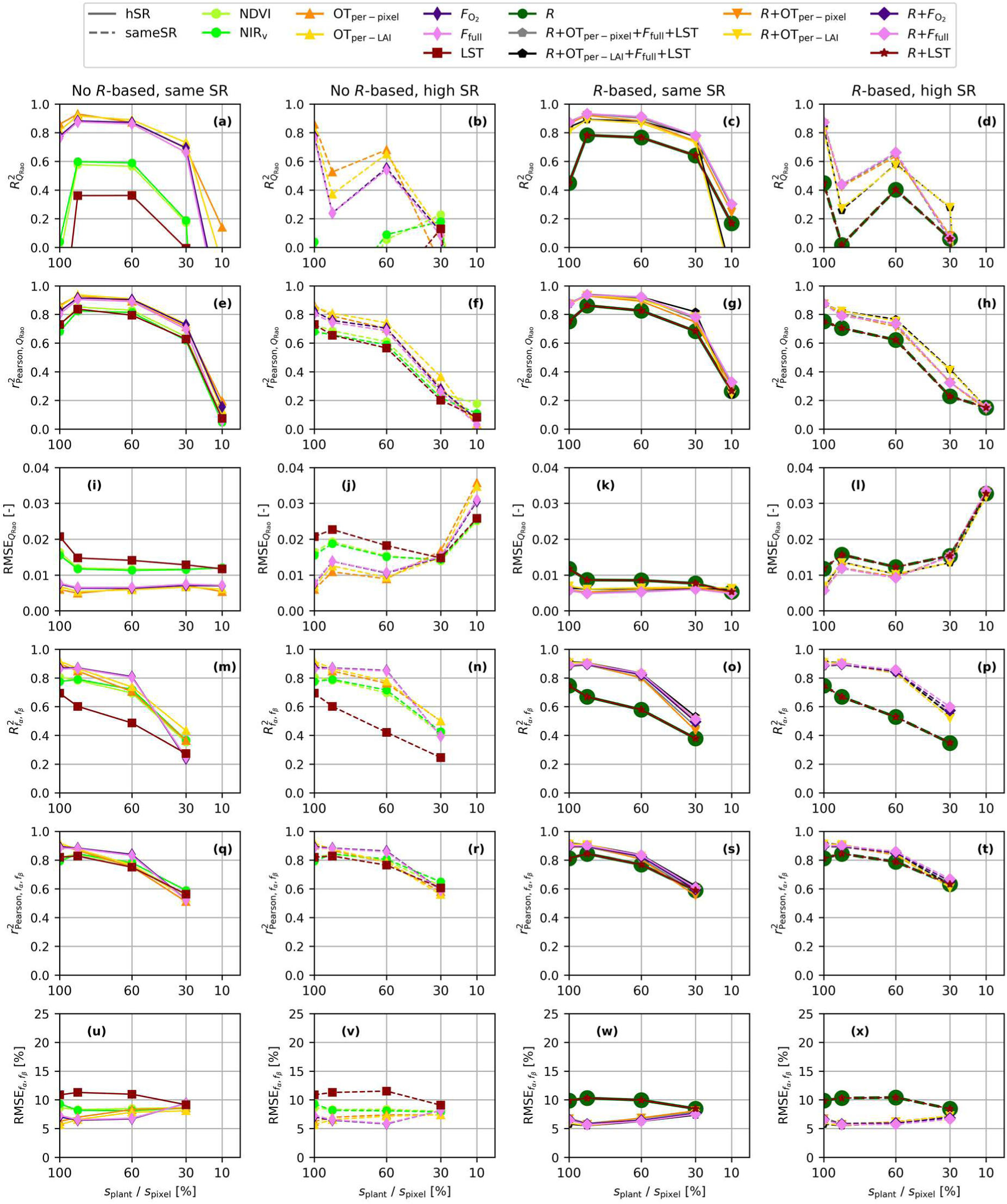
Evaluation of the effect of the spatial resolution (defined as the ratio of the plant to the pixel sizes) on the remote sensing estimation of vegetation *Q*_Rao_ (a-l) and the fractions of alpha or beta-diversity computed via variance-based diversity partitioning (m-x). Coefficient of determination (*R*^2^, a-d, m-p), Pearson correlation (*r*^2^_Pearson_ e-h, q-t), and root mean squared error (RMSE, i-l, u-x). The columns separate the analyses using the full hyperspectral reflectance spectrum between 400-2500 nm (“*R*-based”, columns 3 and 4) from those using only vegetation indices or other signals alone (“No *R*-based”, columns 1 and 2), as well as those cases where the remote sensing estimatesa re compared with field estimates at the same spatial resolution by degrading the field data resolution to meet the remote sensing one (< 100 %, “same SR” columns, 1 and 3) or at the maximum spatial resolution (100 %, “high SR”, columns 2 and 4). Field plant trait diversity metrics are computed using leaf area index as a proxy of abundance (“per-LAI”).

## 4. DISCUSSION

We used BOSSE to answer five fundamental questions regarding the estimation of plant functional diversity from passive optical spectral imagery. Despite inherent limitations, the simulations enabled us to test our hypotheses using consistent data across a wide range of scenarios that encompassed different combinations of plant functional types (determined by the climatic zone), spatial distributions of species, environmental conditions, and the associated vegetation dynamics (local meteorology). Here, we compare the results of BOSSE simulations with those reported for empirical data and provide guidelines for future analyses.

### Q1. How should remote sensing characterize functional diversity in large surfaces (sites)?

BOSSE simulations suggest that averaging the diversity of small samples (e.g., 3-by-3 windows) enables a more accurate estimation of plant functional diversity than large samples (e.g., the whole study area) (H1). This result aligns with previous findings based on simulations lacking the explicit location of species. Pacheco-Labrador et al. (2023) found that variance- and *Q*_Rao_-based *γ*-diversity were less accurately estimated than *α* and *β*-diversity, which was hypothesized to relate to the number of samples. This finding is relevant not only for large-scale diversity monitoring but also for assessing the relationships between biodiversity and ecosystem functions or other processes since: 1) the relationships emerge in part from the interactions between multiple individuals and species (de Bello et al., 2021); and 2) functions and processes are estimated within relatively large areas such as eddy-covariance footprints (Chu et al., 2021), or can be integrated from multiple sampling plots characteristic of the study site.

Research resources limit field sampling, and the challenge is obtaining a sample representative enough of the studied area. For biodiversity, a representative sampling area can be determined using rarefaction curves (Stier et al., 2016; Walker et al., 2008). Conversely, the remote imagers effortlessly and exhaustively sample the Earth’s surface, raising the question of how to compute FDMs (e.g., pixels in moving windows or other grouping criteria, sample sizes). Different studies have examined the effect of the number of pixels or the area used to compute FDMs on their value, finding power-law relationships (Cimoli et al., 2024; Durán et al., 2019; Schneider et al., 2017). However, little attention has been paid to the impact on the relationship between remote sensing and field plant trait FDMs. Tagliabue et al. (2020) found similar correlations between spectral and optical trait-based diversities for 9-by-9 and 3-by-3 pixel moving windows, but the pixel size (1 m) was smaller than the tree crowns imaged, which likely reduced the degree of spectral mixing. Using BOSSE, we further assessed the sample size effect on the relationships between plant trait maps and remote sensing imagery estimates of functional diversity. We found saturating relationships for *Q*_Rao_ and *f_α_* where the estimation performance decreased with increasing window size (Fig. 3, S5-S9).

In this work, we used moving windows to compute FDMs; however, other authors have suggested relying on mapping units of more ecological relevance instead. Laliberté et al. (Laliberté et al., 2020) indicated that windows would not estimate *β*-diversity of the site as a whole. Additionally, Rossi et al. (Rossi et al., 2024) reported that using pasture parcels captured species richness better than previous works. Our results do not oppose the use of mapping units, but rather discourage gathering large numbers of pixels to compute FDMs. Our answer to Q1 shows that *Q_Rao_* from ecologically significant units could be computed as the mean of small subsamples of the patch (“mean” or “median” approach), instead of using all the pixels of the patch together (similar to the “scene” approach). We did not explore the optimal sample size to compute FDMs, but found that estimation performance sometimes maximized at mid-sized windows (Fig. S8-S13). Still, this choice will depend on the spectral variable used and the particular uncertainty of each dataset (Chen et al., 2018). Advances in uncertainty propagation in remote sensing (J. Gorroño et al., 2024) will support these choices.

Finally, we did not evaluate the combined impact of other factors, including degraded spatial resolution, nor did we simulate pixels smaller than the plants. Dealing with the mismatch between the spatial resolution of imagery and variable plant sizes remains an open challenge (Rossi et al., 2021a). Considering our findings and those of Tagliabue et al. (2020), we hypothesize that the spatial data should be integrated to the individual level, when possible.

### Q2. Which sources of spectral information (or combinations thereof) of spectral variables can best capture plant functional diversity at the site scale?

Optical traits (e.g., estimated from reflectance factors) are the most suitable remote sensing variables for estimating plant functional diversity; however, these could be supported by the reflectance factors computed from the “median” of small groups of pixels (H2). BOSSE simulations have ranked OT and *F* as the best estimators of plant functional diversity, likely because these accounted for confounding factors such as bare soil. Moreover, optical traits can be robust to retrieval uncertainty, as equifinal solutions can maintain the variability of plant traits (Pacheco-Labrador et al., 2022). The better performance of *F* over vegetation indices matches former findings in a closed forest (Tagliabue et al., 2020); however, a large variability of performances could be found when evaluations were performed “within sites” (Fig. 4, S3). This result highlights the risk of drawing methodological conclusions from isolated local studies. Spectral indices (NDVI and NIR_v_) exhibited lower performance than *R*, likely because the missing spectral regions in the dataset result in a loss of plant trait information (Pacheco-Labrador et al., 2022). Nonetheless, Gomarasca et al. (2024) found a better representation of ecosystem functional properties using Sentinel-2 NIR_v_ rather than the richer set of spectral bands available, which might be due to accumulated uncertainties, lower spatial resolution of SWIR bands, or the fact that information contained in the index was more relevant to the ecosystem functional properties analyzed. LST (and *R*+LST) resulted in the worst estimator of plant functional diversity, which may be due to the ignored strong soil contributions or a strong dependence on soil and vegetation water fluxes, less related to the plant traits governing spectral light-matter interactions. The performance for the “median” of the small moving windows instead of the “mean” reduced the differences between spectral signals. Then, the performance of *R*, LST, and *R*+LST became comparable to that of the other metrics, which suggests the “median” might alleviate skewed contributions of background effects. Overall, the combination of *R* with other spectral variables (OT, *F*, or LST) did not significantly improve the correlation, accuracy, or performance of the retrievals. This suggests that *R* already integrates a sufficiently large amount of trait information (at least for the traits analyzed), which is not enhanced by the other variables. This is relevant since *F* and LST images are usually noisier and feature coarser spatial resolutions than *R* data (and the derived optical traits). The fact that they are not fundamental for capturing plant functional diversity reduces the requirements on remote sensing data. In practice, *R* and OT would feature the same spatial resolution, and could be used together with a “per-LAI” abundance approach and “median” aggregation to alleviate retrieval and measurement uncertainties. Additionally, whereas considering noise and spatial resolution *F* and LST would provide no added value compared to *R* or OT, they might become more relevant for the analysis of the relationships between biodiversity and ecosystem functions. BOSSE emulators uncertainties scale according to those expected for the different remote sensing data simulated, and OT retrieval can also encompass considerable degrees of error (J. Pacheco-Labrador et al., 2025). However, unlike former studies (Pacheco-Labrador et al., 2022), we did not add random noise to the simulations, which could be assessed in future studies using BOSSE.

### Q3. How should remote sensing estimates be validated/compared with plant functional diversity measurements?

Weighting plant trait FDMs “per-LAI” improved the comparison with remote sensing-based FDMs “per-pixel”, but not in all cases (Fig. 5). For *Q*_Rao_, “per-LAI” enhances accuracy and precision at the expense of correlation strength. In contrast, the benefits of this approach for variance-based diversity partitioning are not as clear. Moreover, the “per-pixel” abundance exhibited lower sensitivity to the spatial scale (window or patch size) of the FDM (Fig. S8-S13). H3 was only partially met.

Overall, the differences between both approaches (“per-LAI” and “per-pixel”) are slight for the spectral variables that perform well to capture diversity (e.g., optical traits), both in performance (Fig. 4) and resistance to the degradation of the spatial resolution (Fig. 7). This might be because LAI, which strongly explains spectral signals’ variance (e.g., Verrelst et al. (2015)), was included as a trait in the computation of the FDMs in the “per-pixel” approach. Overall, the performance is maximized when the metrics from optical and plant traits are computed using the same approach (“per-pixel” or “per-LAI”). The most considerable improvements occur for the least performing variables (e.g., *R*, spectral indices, and LST, Fig. 3-4), which may be because these ignore the presence of bare soil in the window used to compute diversity, and we implemented no method to estimate it.

The extensive sampling of remote sensing imagery raises new methodological questions, such as the identification of bare soil, dead vegetation, and other elements yet to be solved (Cimoli et al., 2024; Pacheco-Labrador et al., 2022; Rossi et al., 2021a). So far, the pixel has most often been proposed as the measure of abundance for computing FDMs from spectral imagery (Laliberté et al., 2020; Rocchini et al., 2017; Rossi et al., 2021b). Alternatively, Tan et al. (Tan et al., 2023) proposed using fractional cover over density. In contrast, as demonstrated in this work, optical trait retrieval provides the opportunity to utilize LAI to weight the contribution of individual pixels to the diversity metric. In fact, LAI is a key variable scaling from leaf to canopy measurements in remote sensing studies (Asner and Martin, 2008). However, ecology studies usually rely on the number of individuals, the area they occupy, their density, or cumulated biomass as a proxy of their abundance when computing diversity metrics (Villéger et al., 2008). The key question yet to be answered is whether taxonomic and biomass-based abundances correspond to the importance of different plants in the signals captured by remote sensing, and whether FDMs optimized for remote sensing are still helpful for ecological analyses. While LAI is a key variable in remote sensing, in ecology, biomass is instead used as an explanatory variable of different processes (Grime, 1998). However, they are to some extent correlated (Goswami et al., 2015; Pandey et al., 2019). Kattenborn et al. (2017) demonstrated that both LAI and specific leaf area exhibit similar explanatory power in discriminating ecological strategies using spectral data, which may help to bridge the gap between the two disciplines. Alternatively, remote sensing could use the same abundance estimates as ecologists. For example, biomass could be directly estimated (Tian et al., 2023). Also, high spatial resolution imagery (spatial resolution ≥ 100 %) could be used to discriminate and count individual plants (Brandt et al., 2020). Furthermore, unmixing methods could estimate the fraction of the pixels covered by different species (Rossi and Gholizadeh, 2023), spectral species (Féret and Asner, 2014). Yet their role in the estimation of plant functional diversity metrics must be further assessed.

### Q4. When (in the phenological year) can remote sensing best capture site-scale plant functional diversity?

Plant functional diversity uncertainty minimizes and remains stable once the mean LAI of the site exceeds 1 m^2^ m^-2^. As hypothesized, BOSSE simulations showed that the RMSE in the retrieval of plant functional diversity does not necessarily minimize at the green peak, but most often at intermediate development stages (Fig. 6, S4). Still, “per-LAI” abundance places RMSE minima at the seasonal peak more often. The dependence of uncertainty on LAI might explain part of the large variability in performance observed in local analyses (“within sites”, Fig. 2, 4, 5). BOSSE simulations caution that diversity estimation is more uncertain when LAI is low, which can compromise the interpretation of functional diversity changes across seasons (Thornley et al., 2022) or the partitioning of the temporal component of diversity (Rossi et al., 2021a). Our work does not assess the information content in phenological diversity; however, BOSSE could also help to test different methodologies to evaluate phenological diversity and their robustness against low LAI or plant coverage, and explore open questions left by previous research (e.g., Girardello et al. (2024)).

### Q5. Which approaches and remote sensing variables are more resistant to the effects of suboptimal spatial resolution?

No spectral variable is fully robust to the degradation of spatial resolution. As expected, the best estimators are also the most resistant (H5.1). We also found that the “per-LAI” weighting slightly improved the resistance of some variables, such as *R* (H5.2), and that under suboptimal resolution, field data can only be reliably compared with remote sensing estimates at the same resolution (H5.3). Even in this case, BOSSE revealed a limitation for pixels more than three times larger than the plants (<30%).

Spatial resolution is the most limiting factor in the remote sensing of vegetation diversity. Most works analyzing the effect of spatial resolution on the estimation of plant diversity have focused on taxonomic diversity (Fassnacht et al., 2022; Gholizadeh et al., 2019; Ludwig et al., 2024; Wang et al., 2018a), likely because of the difficulty of sampling traits at the required extents and resolutions.

Previous non-spatially explicit simulation works showed that spatial resolution strongly reduced the capability of estimating plant functional diversity from spectral data, and that computing plant trait and remote sensing-based diversity metrics at the same spatial resolution (mimicking RS pixels in the field as in Hauser et al. (2021)) improved the robustness of the estimation at suboptimal (<100 %) resolution (Pacheco-Labrador et al., 2022).

While we hypothesized that “per-LAI” abundance would be more robust than “per-pixel”, this improvement was only relevant at the coarsest spatial resolution (10 %), where estimation performance was already low. As foreseen, when remote sensing estimates were compared with high spatial resolution field data (“high SR”), the estimation performance rapidly decreased. Still, the decrease was not monotonic, suggesting that the cause is not the relative pixel size itself, but the degree of mixture, which depends on the colocation between plants and pixels as well. We also found that the variance-based diversity partition is more robust to suboptimal spatial resolution than *Q*_Rao_.

Our results are consistent with previous studies that have reported limitations in characterizing diversity in certain vegetation types or stages where the relative spatial resolution was low (Gholizadeh et al., 2019; Ludwig et al., 2024; Schweiger and Laliberté, 2022). Our findings on the lesser sensitivity of the variance-based partitioning are also consistent with those presented by Schweiger and Laliberté (2022). Spatial resolution is an inherent challenge for studying plant functional diversity from space, as no predefined spatial configuration is suitable for a mixture of plants of different sizes (Laliberté et al., 2020). Even in the ideal case of segmenting plant individuals from very high spatial resolution, it would be challenging and perhaps not feasible for all ecosystems (e.g., grasslands) (Badourdine et al., 2023; Rossi et al., 2021a). Approaches such as degrading high spatial resolution to the average plant size of the Scene (Cimoli et al., 2024) or segmenting individuals from the image could alleviate this problem.

Biodiversity is one of the targets of the new generation of hyperspectral spaceborne sensors (e.g., CHIME (J. Nieke and M. Rast, 2018)). These missions feature pixel sizes of around 30 m, which, according to our results, limits the estimation of plant functional diversity to forests with tree crown diameters of around 10 m, or similar patches of other plants. Thus, it might be necessary to combine this imagery with high spatial resolution multispectral sensors. An open question is the interpretation or value of “same SR” estimates in an ecological context, as the aim is to produce information relevant for ecological analysis. Answering this question may require closer cooperation between ecology and remote sensing disciplines, along with further research assessing the applicability of remote sensing-based diversity estimates to answer ecological questions.

## 5. CONCLUSIONS

We used the Biodiversity Observing System Simulation Experiment (BOSSE) to test simple, yet relevant hypotheses and methodological questions regarding the capabilities of remote sensing to infer plant functional diversity from space. Optical traits offer the best chances to capture plant functional diversity. However, spatial resolution remains the main limitation, as it is not possible to estimate plant functional diversity using pixels that are more than three times larger than the plants themselves. Remote sensing estimates of plant diversity in large areas should rely on small windows or patches of pixels whose values are averaged. The leaf area index can be a suitable proxy for vegetation abundance. Nonetheless, the estimation becomes more uncertain when soil contribution is salient (e.g., during the dry season).

Despite the inherent limitations of synthetic data, BOSSE has proven to be a useful framework for addressing questions that the lack of sufficient systematic and comparable observations prevents from being tackled. In fact, the simulations revealed a wide variability in performances for local analyses, which suggests caution when interpreting empirical results that are local in nature. We expect BOSSE to support advances in the field of remote sensing of plant functional diversity, and guide necessary tests to be carried out from field measurements and remote sensing-designed validation schemes.

## ACKNOWLEDGEMENTS

JPL acknowledges the project “Integrated Observing Systems and Simulation Experiments to Analyze Biodiversity-Ecosystem Function Relationships in Savanna Ecosystems” PID2023-151046NB-I00 funded by MCIU/ AEI / 10.13039/501100011033 / FEDER, UE) and the ESA Living Planet Fellowship IRS4BEF (ESA Contract No. 4000140028/22/I-DT-lr).

## Declaration of AI and AI-assisted technologies in the writing process

This manuscript’s readability and language correctness were supported by Grammarly. After using this tool, the authors reviewed and edited the content as needed and take full responsibility for the content of the publication.

## CRediT authorship contribution statement

**Javier Pacheco-Labrador:** Writing–original draft, Visualization, Methodology, Investigation, Formal analysis, Data curation, Conceptualization, Funding acquisition. **Ulisse Gomarasca:** Writing– review & editing, Conceptualization. **Daniel E. Pabon-Moreno:** Writing–review & editing, Methodology, Conceptualization. **Wantong Li:** Writing–review & editing, Data curation, Conceptualization. **Mirco Migliavacca:** Writing–review & editing, Conceptualization. **Martin Jung:** Writing–review & editing, Conceptualization, Funding acquisition. **Gregory Duveiller:** Writing– review & editing, Conceptualization, Funding acquisition.

## Data availability

The Python code used to run the simulations and figures of this manuscript can be found in Zenodo: https://doi.org/10.5281/zenodo.17414765. The Biodiversity Observing System Simulation Experiment (BOSSE v1.0) code can be found on GitHub: https://github.com/JavierPachecoLabrador/pyBOSSE; the additional ERA5Land Meteorological time series are available in Zenodo: https://doi.org/10.5281/zenodo.14717038. The pyGNDiv package used to compute functional diversity metrics can be found on GitHub: https://github.com/JavierPachecoLabrador/pyGNDiv-master.

