## Supplementary Material for "Benchmarking remote sensing methods to capture plant functional diversity from space"

#### Figure S1

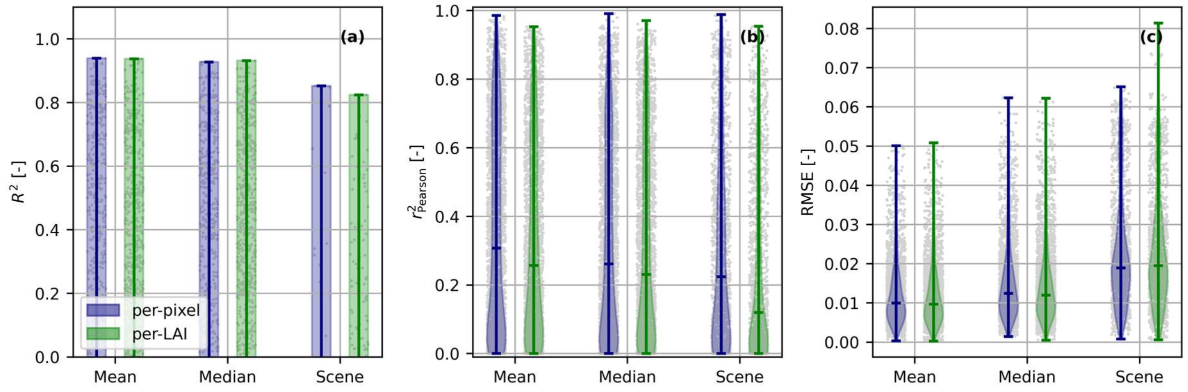


**Figure S1.** Distribution of the statistics resulting from the comparison of functional diversity metrics computed from plant traits and several combinations of remote sensing variables. Plant traits' abundance is represented by different proxies: pixels covered ("per-pixel") or leaf area index ("per-LAI"). Coefficient of determination (a), Pearson correlation coefficient (b), and root mean squared error (c). The comparison is performed "within sites" by comparing the Rao's quadratic entropy index (*Q*) computed at each time stamp of the simulation, site by site.

### Figure S2

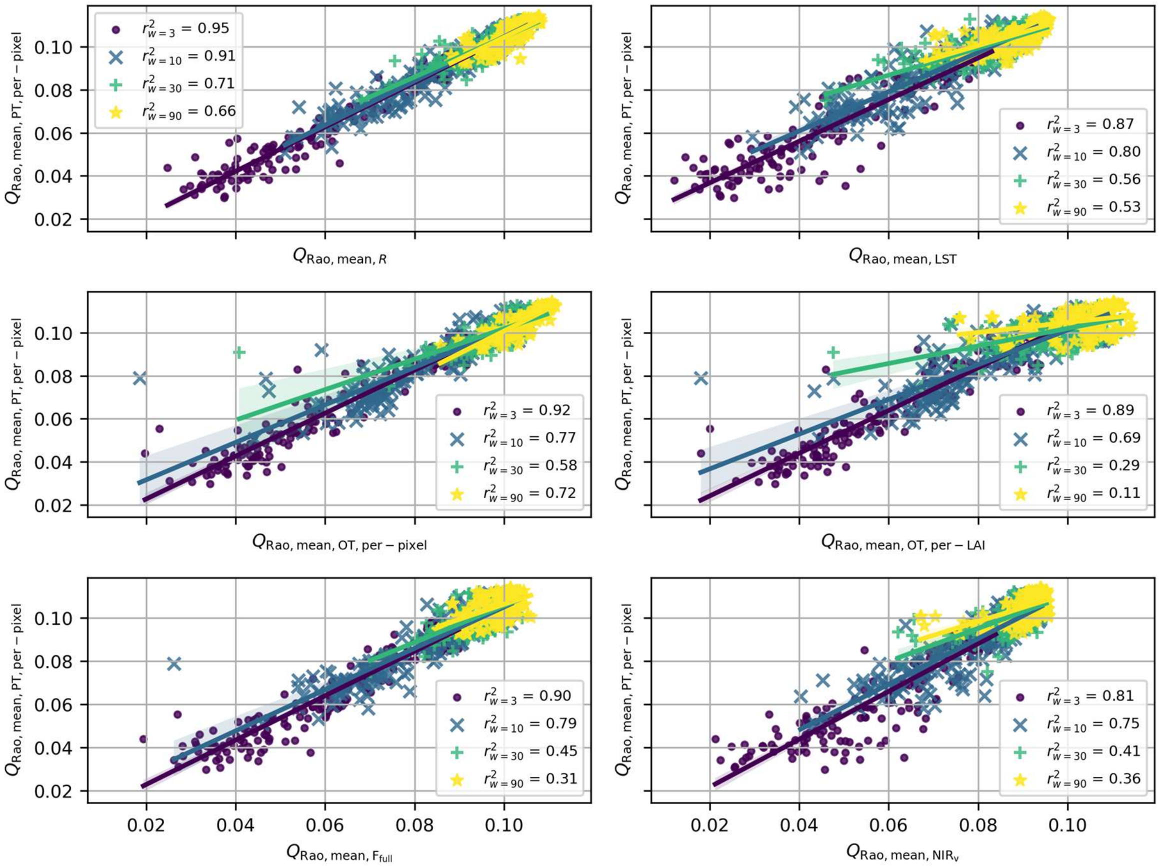


**Figure S2.** Rao's quadratic entropy index ($Q_{\mathrm{Rao}}$) computed from plant traits ($y$-axis) and remote sensing variables ($x$-axis) using windows of different sizes (3, 10, 30, and 90 pixels). The remote sensing variables are hyperspectral reflectance factors ($R$), land surface temperature (LST), optical traits where relative abundance is computed as a function of pixel area (OT, per-pixel), optical traits where relative abundance is computed as a function of leaf area index (OT, per-LAI), hyperspectral sun-induced chlorophyll fluorescence ($F_{\mathrm{full}}$), and near infrared of vegetation ($\mathrm{NIR_v}$).

#### Figure S3

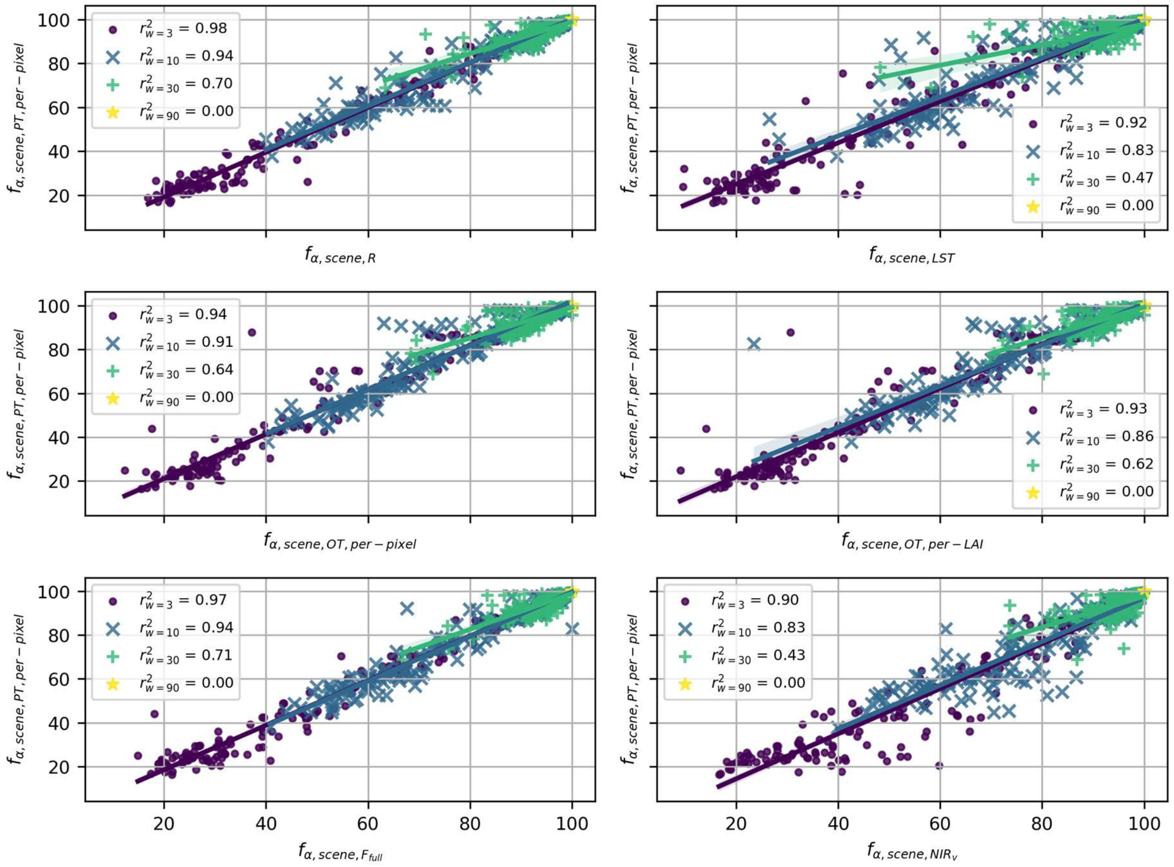


**Figure S3.** Fraction of $\alpha$-diversity ($f_\alpha$) computed from plant traits ($y$-axis) and remote sensing variables ($x$-axis) using windows of different sizes (3, 10, 30, and 90 pixels). The remote sensing variables are hyperspectral reflectance factors ($R$), land surface temperature (LST), optical traits where relative abundance is computed as a function of pixel area (OT, per-pixel), optical traits where relative abundance is computed as a function of leaf area index (OT, per-LAI), hyperspectral sun-induced chlorophyll fluorescence ($F_{\text{full}}$), and near infrared of vegetation ($NIR_v$).

### Figure S4

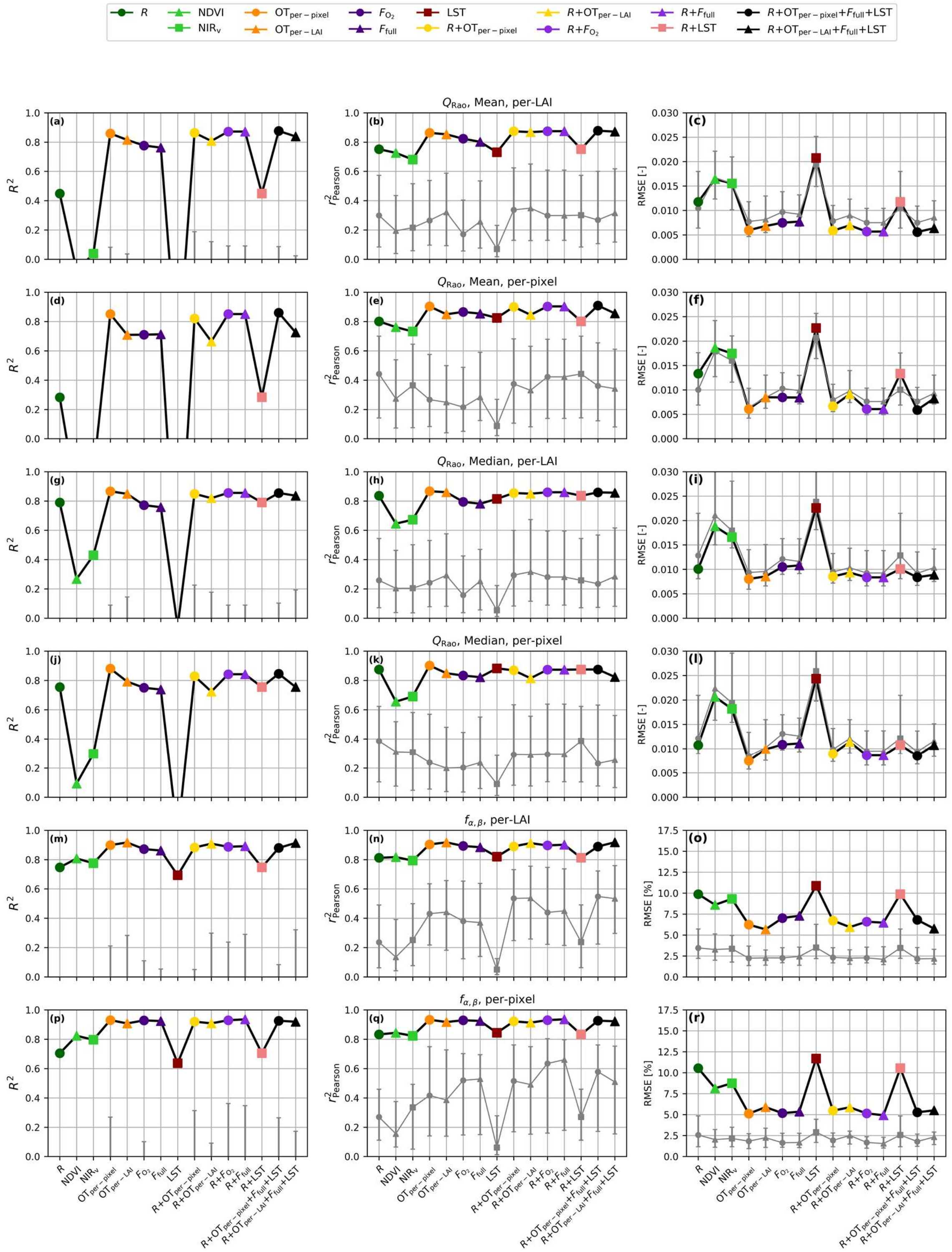


**Figure S4.** Statistics resulting from the comparison of functional diversity metrics computed from plant traits and several combinations of remote sensing variables computed from the the mean of small moving windows for $Q_{\mathrm{Rao}}$ using LAI (a-c) or pixel coverage (d-f) as a proxy of abundance, from median of small moving windows for $Q_{\mathrm{Rao}}$ using LAI (g-i) or pixel coverage (j-l) as a proxy of abundance, and the fraction of alpha and beta-diversity using LAI (m-o) or pixel (p-r) as a proxy of abundance. Coefficient of determination (a, d, g, j, m, p), Pearson correlation coefficient (b, e, h, k, n, q), and root mean squared error (c, f, i, l, o,r).

### Figure S5

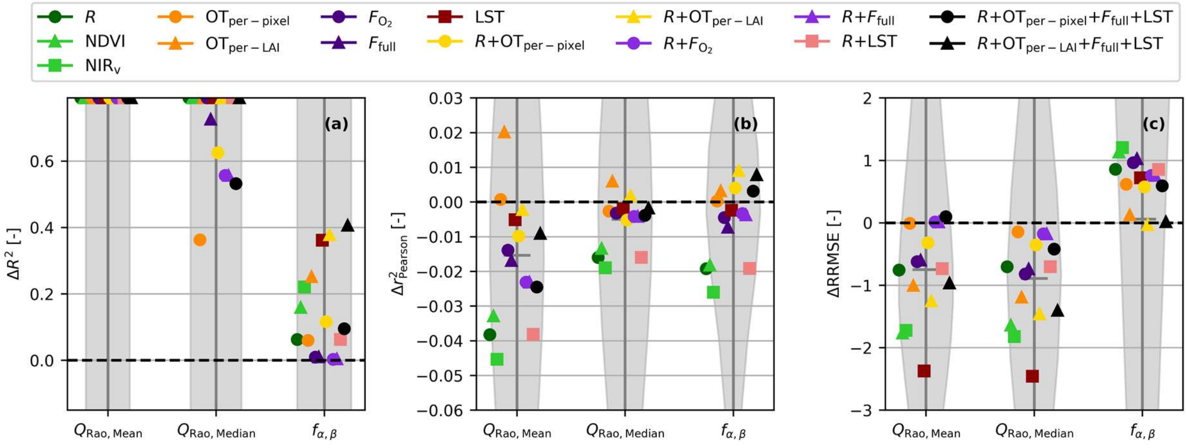


**Figure S5.** Difference between the statistics resulting from the comparison of remote sensing-based and field-based functional diversity metrics. The difference (*Δ*) is built as the difference of the statistics for the case where field-based abundance is represented by the leaf area index ("per-LAI") minus the case where it is represented by the pixel coverage ("per-pixel"). Coefficient of determination (a), Pearson correlation coefficient (b), and root mean squared error (c). The statistics are truncated to the plot axis for clarity, and represent a comparison of the time series of diversity metrics for each Scene simulated individually ("within sites").

### Figure S6

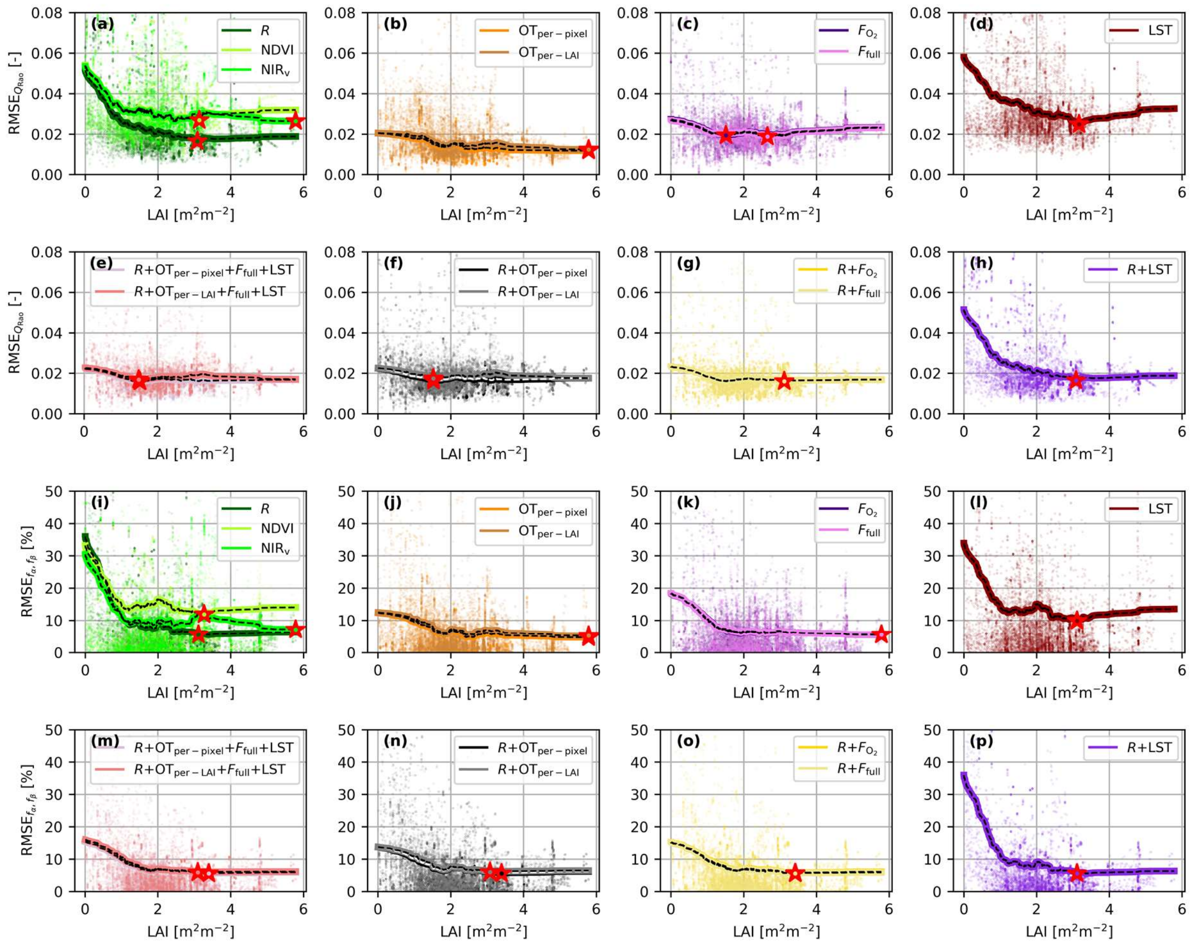

**Figure S6.** Relationship between mean scene leaf area index (LAI) and the root-mean-squared error of the plant functional diversity (Rao's quadratic entropy index ($Q_{\text{Rao}}$)) and the fractions of alpha and beta-diversity (combined) at each simulation timestamp. Field-based functional diversity is computed using pixel area for abundance ("per-pixel"), and it is estimated with different remote sensing sets of variables: reflectance factors ($R$), optical traits used to calculate diversity using the pixel area ($R$+$\text{OT}_{\text{per-pix}}$) or leaf area index ($R$+$\text{OT}_{\text{per-LAI}}$) for abundance, sun-induced chlorophyll fluorescence radiance retrieved at the $O_2$ absorption bands ($F_{O_2}$), the entire emission spectrum ($F_{\text{full}}$), land surface temperature (LST), and combinations of the former ones.

### Figure S7

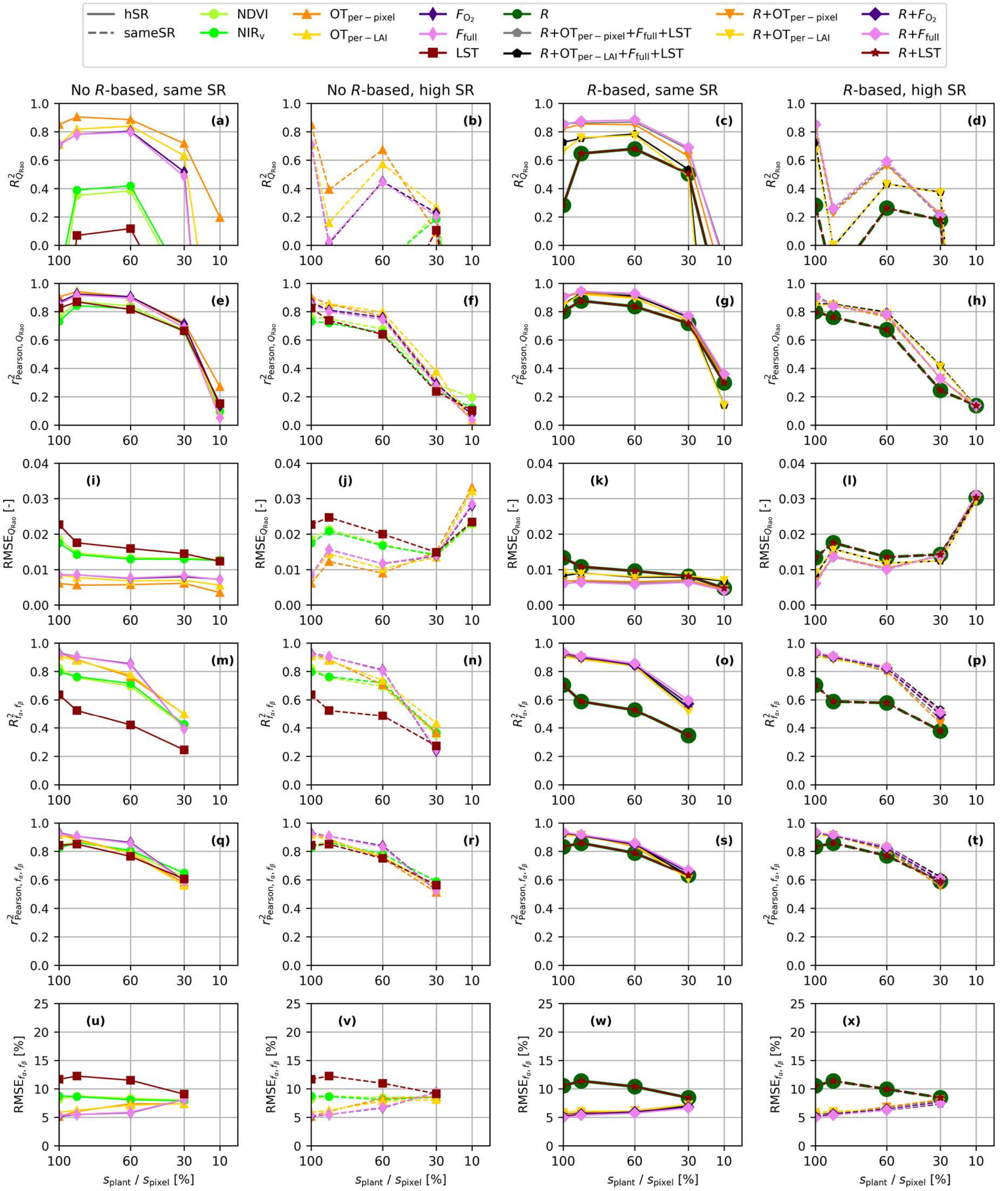


**Figure S7.** Evaluation of the effect of the spatial resolution (defined as the ratio of the plant to the pixel sizes) on the remote sensing estimation of vegetation $Q_{\mathrm{Rao}}$ (a-l) and the fractions of alpha or beta-diversity computed via variance-based diversity partitioning (m-x). Coefficient of determination ($R^2$, a-d, m-p), Pearson correlation ($r^2_{\mathrm{Pearson}}$ e-h, q-t), and root mean squared error (RMSE, i-l, u-x). The columns separate the analyses using the full hyperspectral reflectance spectrum between 400-2500 nm ("$R$-based", columns 3 and 4) from those using only vegetation indices or other signals alone ("No $R$-based", columns 1 and 2), as well as those cases where the remote sensing estimatesa re compared with field estimates at the same spatial resolution by degrading the field data resolution to meet the remote sensing one (< 100 %, "same SR" columns, 1 and 3) or at the maximum spatial resolution (100 %, "high SR", columns 2 and 4). Field plant trait diversity metrics are computed using the pixel area as a proxy of abundance ("per-pixel").

**Figure S8**

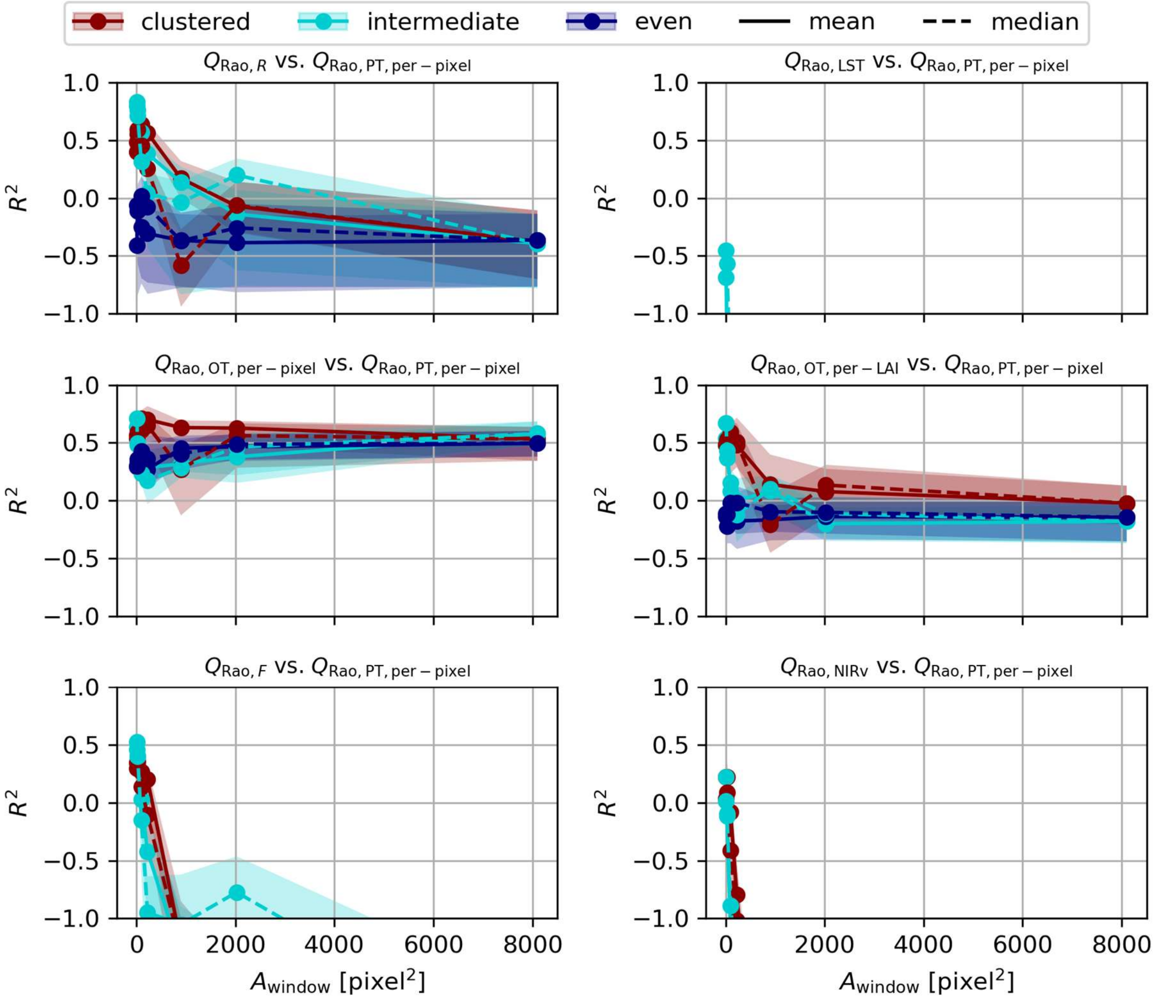


**Figure S8.** Coefficient of determination of the relationship between plant trait-based functional diversity ($Q_{Rao}$, "per-pixel") vs. remote-sensing-based functional diversity ($Q_{Rao}$) separated by vegetation distribution type as a function of the moving window area.

#### Figure S9

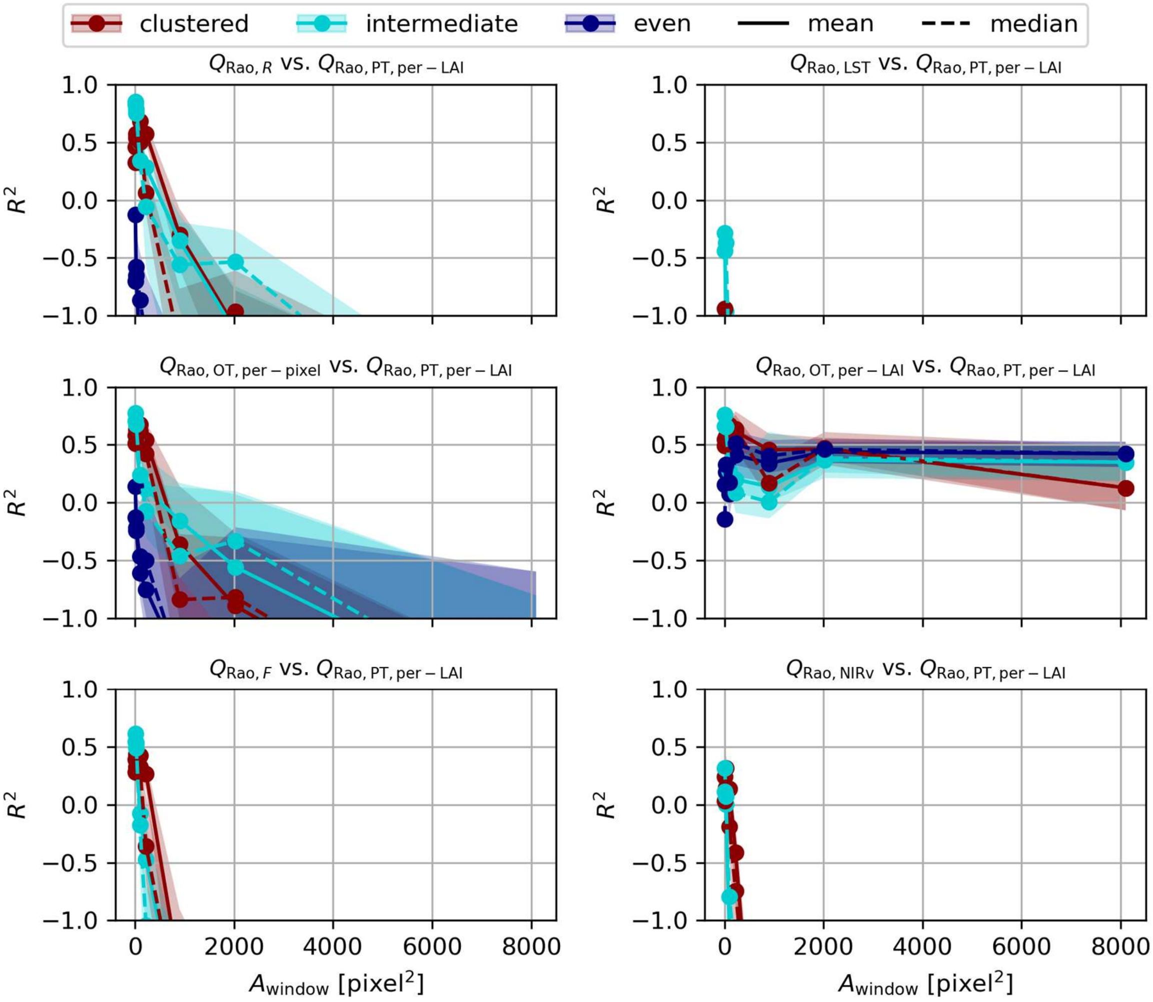


**Figure S9.** Coefficient of determination of the relationship between plant trait-based functional diversity ($Q_{Rao}$, “per-LAI”) vs. remote-sensing-based functional diversity ($Q_{Rao}$) separated by vegetation distribution type as a function of the moving window area.

**Figure S10**

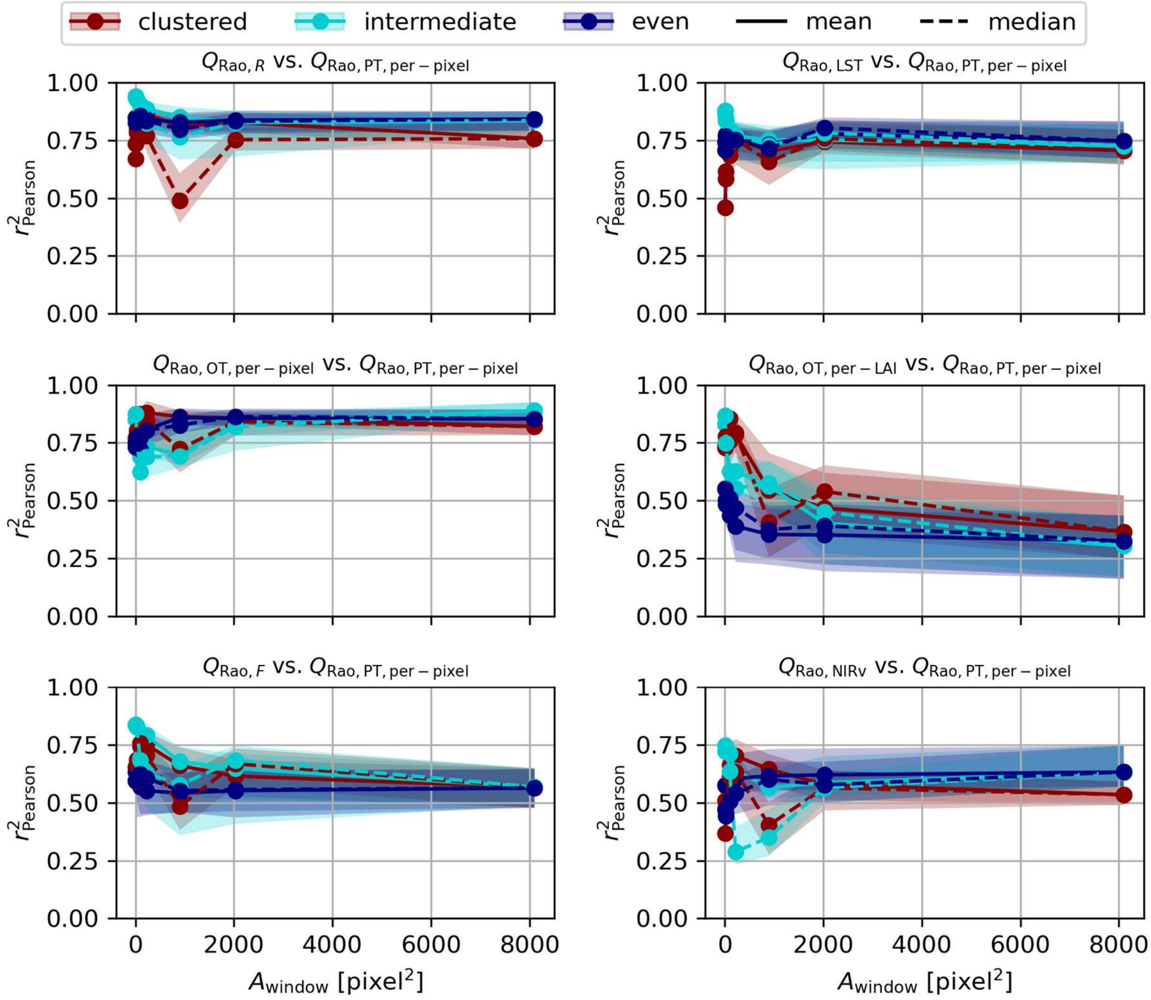


**Figure S10.** Pearson correlation coefficient of the relationship between plant trait-based functional diversity ($Q_{Rao}$, "per-pixel") vs. remote-sensing-based functional diversity ($Q_{Rao}$) separated by vegetation distribution type as a function of the moving window area.

#### Figure S11

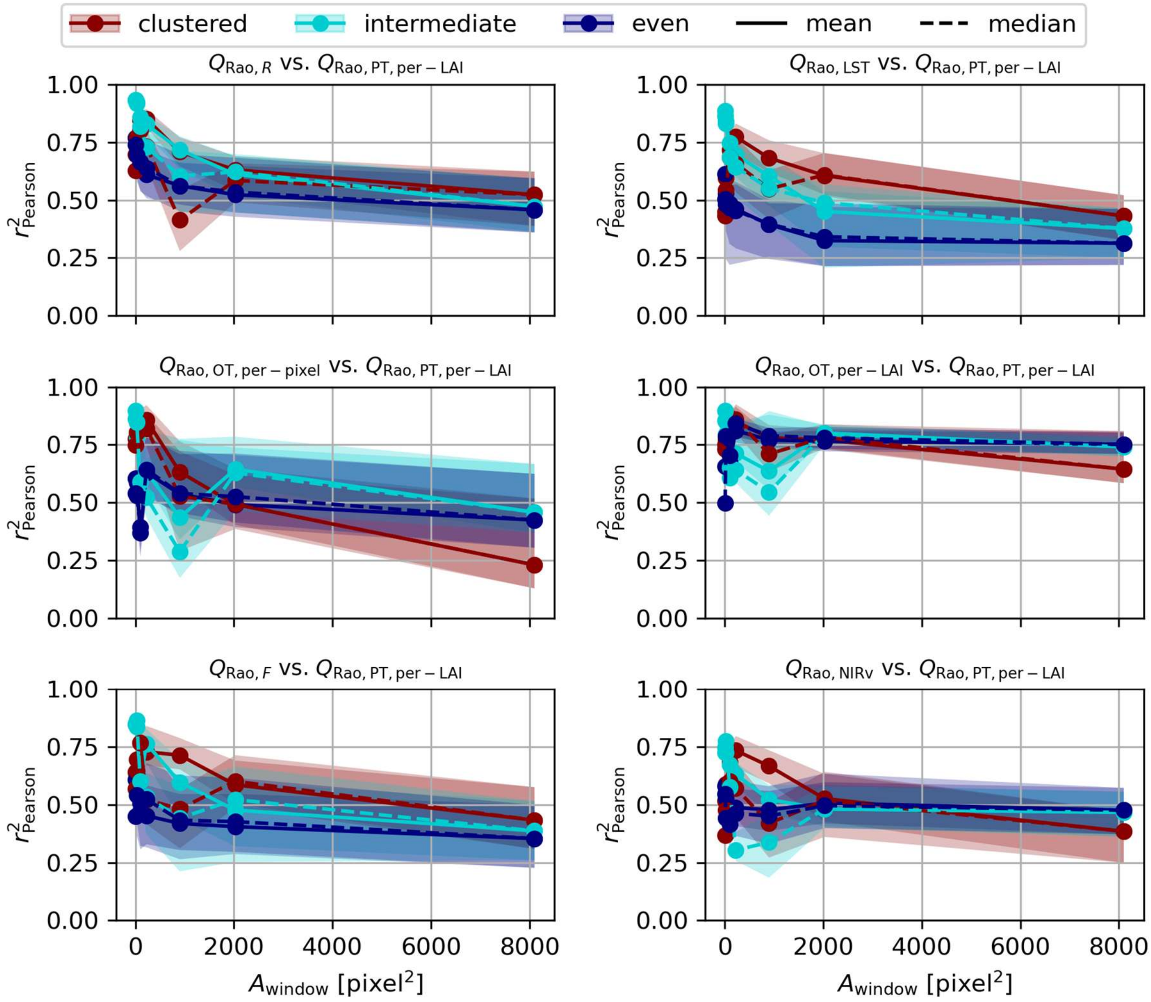


**Figure S11.** Pearson correlation coefficient of the relationship between plant trait-based functional diversity ($Q_{Rao}$, "per-LAI") vs. remote-sensing-based functional diversity ($Q_{Rao}$) separated by vegetation distribution type as a function of the moving window area.

**Figure S12**

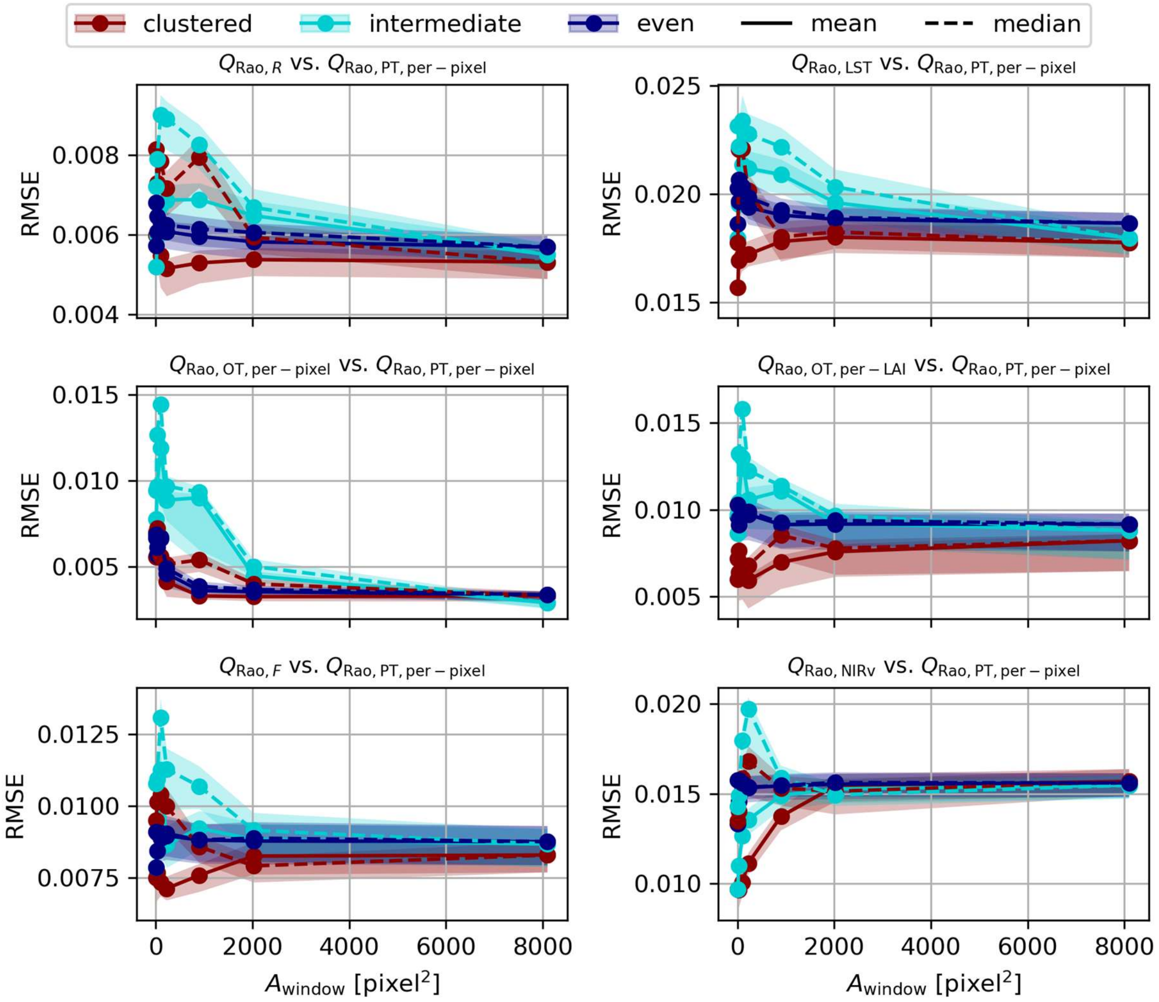


**Figure S12.** Root mean squared error of the relationship between plant trait-based functional diversity ($Q_{\mathrm{Rao}}$, "per-pixel") vs. remote-sensing-based functional diversity ($Q_{\mathrm{Rao}}$) separated by vegetation distribution type as a function of the moving window area.

#### Figure S13

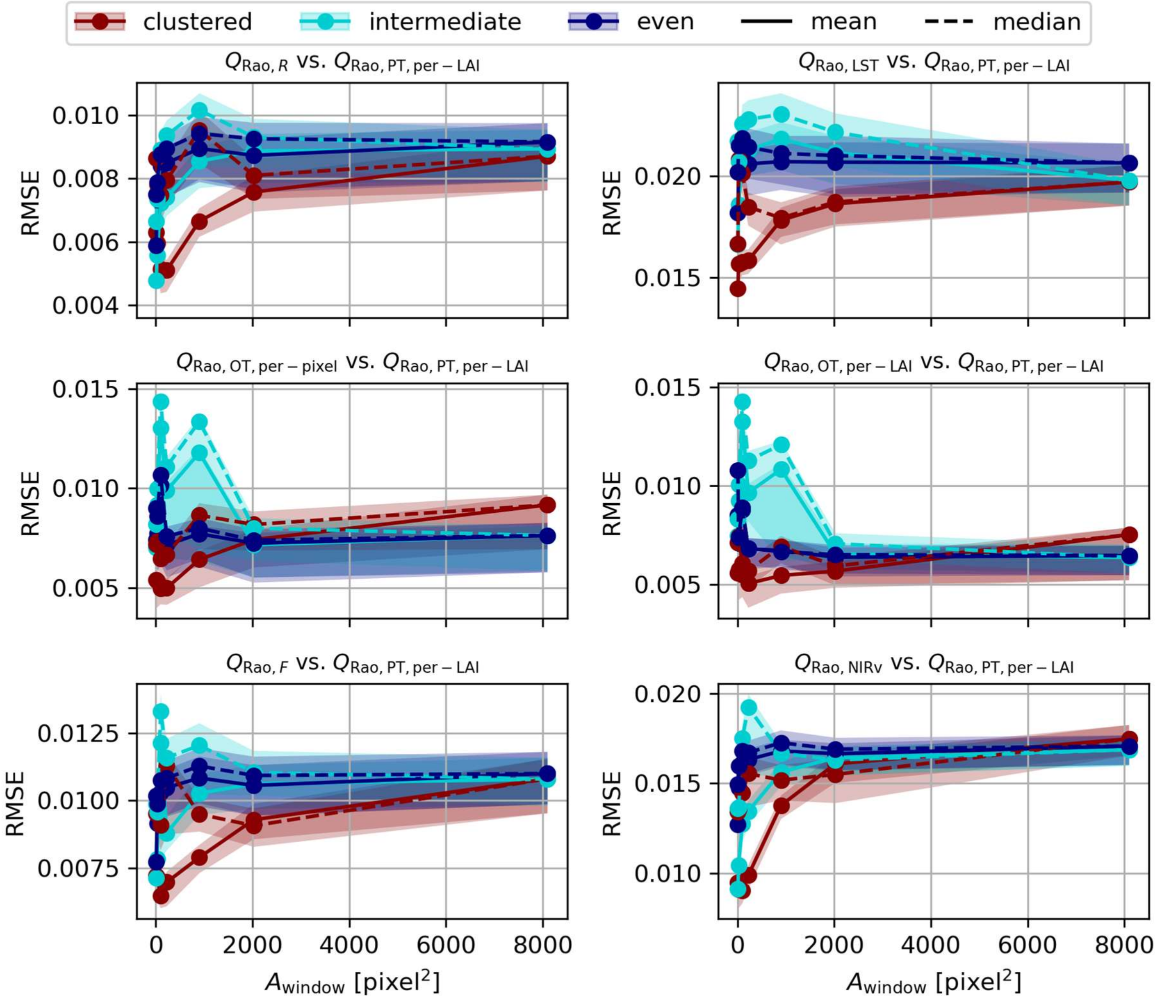


---

**Figure S13.** Root mean squared error of the relationship between plant trait-based functional diversity ($Q_{\mathrm{Rao}}$, "per-LAI") vs. remote-sensing-based functional diversity ($Q_{\mathrm{Rao}}$) separated by vegetation distribution type as a function of the moving window area.
